# Epithelial to mesenchymal transition and M2 macrophages drive aggressiveness in murine and human sarcomatoid urothelial carcinoma

**DOI:** 10.64898/2026.09.30.755727

**Authors:** Yujiro Hayashi, Roy Elias, Eugene Douglass, Akinaru Yamamoto, Matthew Schuler, Mingxiao Feng, Ekatherina Y. Batourina, Anne E. Geller, Woonyoung Choi, Gabriela Colocho, Alyssa Arbuiso, Samuel Jin, Adam K. Aragaki, Charles Ruland, Stanley Rapiey, Jean Hoffman-Censits, Max Kates, Sunil H. Patel, Nirmish Singla, Armine Smith, Stephanie Russell, Huili Li, Joshua I. Warrick, Ezra Baraban, Karin Allenspach, Cathy L. Mendelsohn, Andres Matoso, Noah M. Hahn, David J. McConkey, Burles A. Johnson

**Affiliations:** Greenberg Bladder Cancer Institute, Johns Hopkins University; Department of Oncology, Johns Hopkins University; Department of Urology, Johns Hopkins University; Department of Pathology, Johns Hopkins University; College of Pharmacy, University of Georgia; College of Veterinary Medicine, University of Georgia; Department of Pathology, Columbia University; Department of Pathology, Yale University.

## Abstract

Patients with majority variant subtype histology including sarcomatoid urothelial carcinoma (SUC) have poor outcomes and limited treatment options, and there are few preclinical resources to investigate rare histologic subtypes. We developed three novel variant subtype histology bladder cancer (BC) cell lines by finding that basal BC tumor cells derived from mice treated with N-butyl-N-(4-hydroxybutyl)-nitrosamine (BBN) were initially not competent to grow in immune competent mice, but then acquired tumorigenicity following passaging (“recycling”) in mice lacking mature B and T cells. Three genitourinary oncology pathologists identified BBN966 as SUC, BBN964 as a hybrid between SUC and squamous, and BBN975 as squamous, and each had unique growth characteristics. Using a publicly available dataset and our spatial transcriptomics analysis of a cohort of human tumors with mixed SUC and UC as comparators, we found that consistent with human SUC, BBN966 lack expression of basal and luminal genes. Consistent with human SUC and spatial transcriptomics analysis of canine SUC, BBN966 had the highest epithelial to mesenchymal transition (EMT) upregulation, and all three recycled tumor cells had increased EMT pathway expression in recycled tumor cells. Moreover, increased expression of genes upregulated in all three recycled tumor cells associated with worse overall survival in patients with muscle invasive UC. Similar to human SUC, BBN966 tumors had higher proportion of M2 macrophage gene expression. Finally, we found in the largest cohort of PD-L1 IHC to date that 20 of 38 (52.6%) human SUC had CPS ≥ 50, which was driven mostly by tumor proportion score. Consistent with this, we found anti-PD1 immune checkpoint inhibitor (ICI) significantly inhibited BBN966 tumor growth, while the hybrid BBN964 tumors had mixed response to ICI. Thus, the murine BBN966 tumors exhibited similarities to human SUC, and can be used to examine novel therapies for translation into future SUC clinical studies.

## Introduction

Variant histologic subtypes of urothelial carcinoma (UC), including sarcomatoid urothelial carcinoma (SUC) represent a subset of rare and aggressive bladder cancer with poor overall survival (OS)^1,2^. While more recent reports indicate that chemotherapy^3^ and immune checkpoint inhibitor (ICI) therapy^4^ have potential efficacy in treating SUC, patients with SUC have poor overall survival (OS) when compared with patients with conventional UC^5,6^. Moreover, practice-changing clinical trials in conventional UC did not include patients with majority variant histologic subtypes including SUC^7–10^, and real-world data suggest that therapy such as enfortumab vedotin for patients with conventional UC is not as efficacious when used in patients with majority variant histologic subtypes^11^. Thus, novel therapeutic strategies are urgently needed for patients with SUC.

One challenge to therapeutic development in SUC (and other variant histologic subtypes) is the relative lack of models that faithfully replicate human disease. Current available models include a patient derived xenograft^12^ and patient derived organoids^13^. The BBN [N-butyl-N-(4-hydroxybutyl) nitrosamine] model of carcinogenesis yields SUC in a minority of mice^14^, however inability to predict which mice will develop this phenotype makes therapeutic intervention studies of SUC using this model difficult. There is not a mouse model of SUC available that (1) is syngeneic and therefore permits study in a fully immunocompetent host, allowing for optimized study of immunotherapeutics, and (2) has been documented to replicate the tumor microenvironment (TME) of human SUC. Here we integrate murine with human and canine SUC data to describe the generation of an optimized mouse SUC cell line, a cell line that is a hybrid between SUC and squamous UC, and a squamous UC cell line, each with unique growth kinetics and phenotypic characteristics. We demonstrate that the SUC cell line BBN966 replicates the phenotypic characteristics of human SUC, and when implanted, exhibits similarities with the human SUC TME, such as epithelial to mesenchymal transition (EMT), and unlike the BBN964 cell line, modest response to immune checkpoint inhibitor (ICI) therapy, and predominance of macrophages with an immune suppressive phenotype. Thus, the BBN966 model represents a novel tool for use to improve therapeutic development in SUC, and can be used in conjunction with the BBN964 and BBN975 cell lines to study differences between SUC and UC.

## Materials and Methods

### Mice

All mouse experiments were approved by the Johns Hopkins Institutional Animal Care and Use Committee (Protocol Numbers: MO24M110 and MO25M378). All mice used for experiments were C57BL/6J mice purchased from Jackson Laboratories (Cat. No. 000664). Rag1-KO mice were initially purchased from Jackson Laboratories (Cat. No. 002216) and then later bred in our laboratory. All mice used in generation of Fig. 1 were female age matched mice greater than 8 weeks but less than 6 months old.

**Figure 1.**
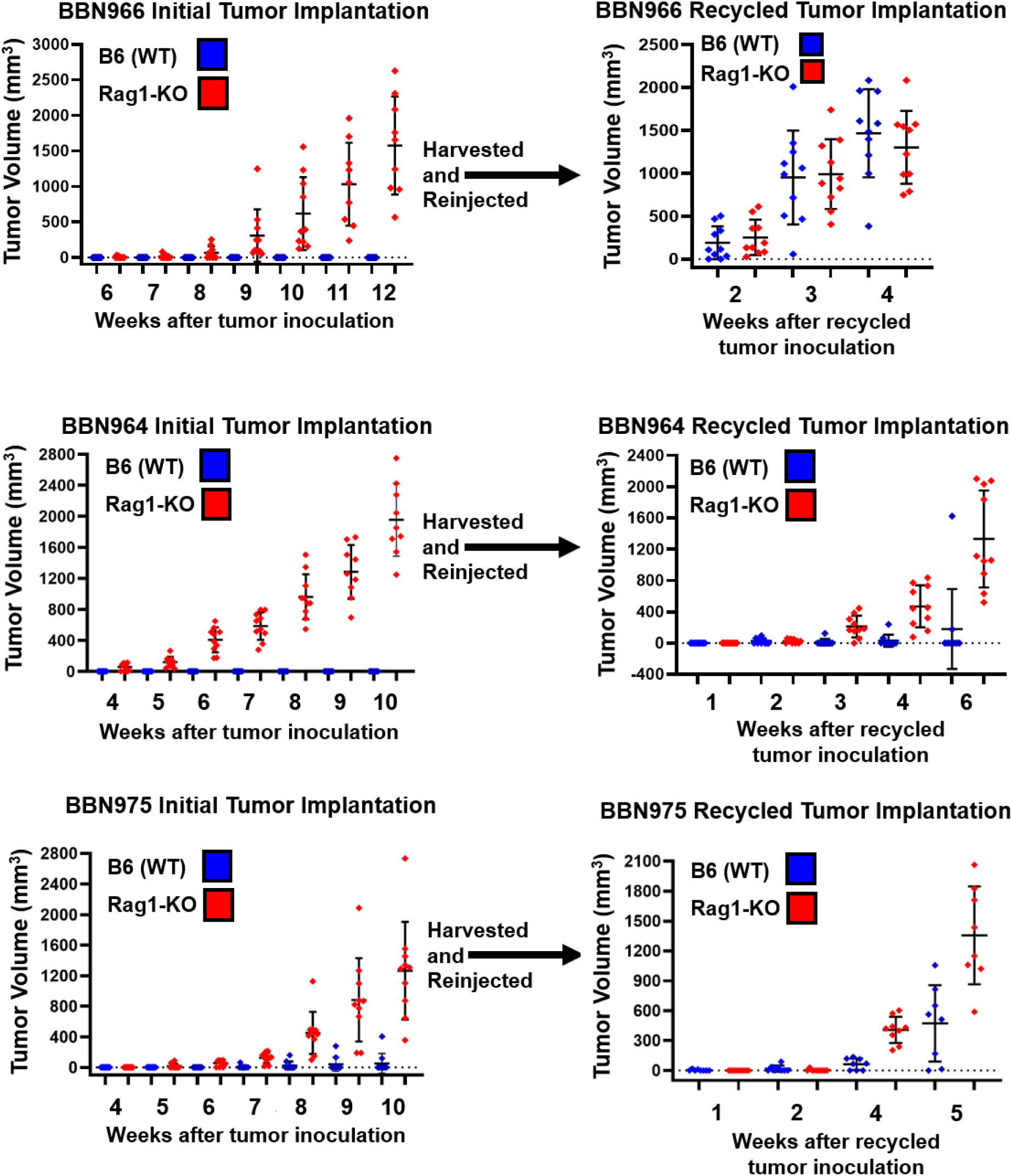
Adaptive immunity inhibits growth of the basal subtype of urothelial carcinoma. BBN966 (top), BBN964 (middle), and BBN975 (bottom) were implanted (5x10^6, s/q) into B6 and Rag1-KO mice (left). At time of harvest, single cell suspensions from tumors grown in Rag1-KO mice from each group were subsequently implanted into naïve B6 and Rag1-KO mice (right). Tumor volumes were measured via calipers.

### Murine RNA extraction and library preparation

Tumor cells were cultured as monolayers in 6-well plates. Total RNA was extracted using the mirVana miRNA Isolation Kit (Thermo Fisher Scientific, Waltham, MA, USA) according to the manufacturer’s instructions. Briefly, cells were washed with phosphate-buffered saline and lysed directly in the culture wells. Extracted RNA was treated with the TURBO DNA-free Kit (Thermo Fisher Scientific) to remove residual genomic DNA according to the manufacturer’s instructions.

RNA concentration was quantified using a Qubit fluorometer (Thermo Fisher Scientific), purity was assessed using a NanoDrop spectrophotometer (Thermo Fisher Scientific), and RNA integrity was evaluated using a TapeStation system (Agilent Technologies, Santa Clara, CA, USA).

RNA-sequencing libraries were prepared from 10 ng of DNase-treated total RNA using the Ion AmpliSeq Transcriptome Mouse Gene Expression Kit (Thermo Fisher Scientific) according to the manufacturer’s instructions. Briefly, RNA was reverse-transcribed using the SuperScript VILO cDNA Synthesis Kit, followed by multiplex amplification with the Ion AmpliSeq Transcriptome Mouse Gene Expression Core Panel. Primer sequences were partially digested, and Ion Xpress Barcode Adapters (Thermo Fisher Scientific) were ligated to the resulting amplicons. The libraries were subsequently purified using AMPure XP beads. The quality of the purified libraries was assessed using a TapeStation system (Agilent Technologies) prior to template preparation and sequencing.

### Immunofluorescence Staining and Analysis

Fourteen paraffin-embedded tumors were sent to the Mendelsohn Laboratory at Columbia University for immunofluorescence staining and image acquisition. Images were collected from 208 tissue sections, with 1–3 unique images captured per slide at 10X and 20X magnification. Image filenames included the block ID, stain and corresponding fluorophore, magnification, and tissue section. Tumor blocks were matched to their corresponding cell line and pathological diagnosis determined by Drs. Andres Matoso, Ezra Baraban, and Joshua Warrick. A Microsoft PowerPoint presentation was created for each tumor block to facilitate review of all images, including the block name, tissue section, magnification, cell line, pathology diagnosis, and fluorophore labels. Dr. Burles Johnson and Matthew Schuler independently reviewed each DAPI image to assess tissue quality, and images with less than 80% cellular coverage or evidence of scanning artifact were excluded from analysis. Of 982 total images, 111 (11.3%) were excluded, leaving 871 images for quantitative analysis. Fluorescence intensity was quantified across the entire imaging field using Fiji (ImageJ) with a custom macro to measure raw integrated density. Red, green, and blue fluorescence channels were processed using the “8-bit Color” option, whereas E-cadherin fluorescence, acquired as a white/gray channel, was processed using the “8-bit” option. All other image-processing and quantification steps were identical across fluorescence channels. Quantitative data were organized by stain and cell line in Microsoft Excel and analyzed using GraphPad Prism. Differences between groups were assessed using one-way analysis of variance (ANOVA) with Tukey’s multiple-comparisons test, with statistical significance defined as p < 0.05.

### Human GeoMx data

Spatial transcriptomics data of the Hopkins cohort of tumors with mixed SUC and UC were generated as previously described^15^.

#### Data processing

Target-level count matrices, ROI annotations, and probe annotations were exported from the GeoMx DSP Data Analysis suite and imported into R using readGeoMx from the standR package^16^, producing a SpatialExperiment object with negative probes identified by the “Negative Probe” code class. Per-ROI quality metrics were computed with addPerROIQC, which also removes genes flagged as undetected across the cohort. ROIs with fewer than 150 nuclei were excluded. Counts were normalized by the trimmed mean of M-values (TMM) method using geomxNorm. Principal component analysis showed no separation by slide, and no batch correction was applied. Analysis was restricted to ROIs annotated as urothelial carcinoma. Each ROI carried a histologic annotation of sarcomatoid or conventional epithelial assigned by a pathologist. <u>*Differential expression.*</u> Normalized objects were converted to DGEList objects with SE2DGEList. Lowly expressed genes were removed with filterByExpr. Dispersion was estimated with estimateDisp (robust), precision weights computed with voom, and linear models fit with lmFit followed by empirical Bayes moderation. The design was a means model on histologic annotation (∼ 0 + tissue_type) and the contrast tested was sarcomatoid minus conventional epithelial. Genes with a Benjamini-Hochberg adjusted p-value below 0.05 were called differentially expressed.

### Canine GeoMx data

#### Data Generation

Spatial profiling of canine urothelial carcinoma biopsy samples was performed using the GeoMx™ Digital Spatial Profiler (DSP; Bruker Spatial Biology, Seattle, WA) through the Flow Cytometry and Immune Monitoring Shared Resource at the University of Arizona. All stainings were performed following steps outlined in the GeoMx™ DSP Automated Slide Prep User Manual (MAN-10151) using the Leica Bond RXm (Leica Biosystems, Deer Park, IL).

Formalin fixed paraffin-embedded sections of canine tissue were cut at 5µm and mounted on Leica Apex BOND Superior Adhesive Slides (cat 3800040, Leica Biosystems, Deer Park, IL). Slides were then deparaffinized, treated to heat-induce epitope retrieval (HIER) for 20 minutes at 100°C, before being treated with 1 µg/mL proteinase K for 15 minutes in order to improve binding and reactivity for hybridization and staining. The tissue sections where then hybridized with the GeoMx™ Cancer Atlas Canine RNA Probes for NGS Compatible w/ Illumina Systems (cat. 121400107, Bruker Spatial Biology, Seattle, WA) overnight.

Following hybridization, the slides underwent a series of stringent washes to remove off-target probes before morphology marker antibody staining with GeoMx™ Solid Tumor TME Morphology Kit Human RNA Compatible (cat. 121300310, Bruker Spatial Biology, Seattle, WA), consisting of antibodies against Pan-CK. Slides were additionally labeled with a custom morphology marker – Vimentin (E-5) Alexa Fluor 647 (cat. SC-373717 AF647, Santa Cruz Biotechnology, Dallas, TX) at a concentration of 10 µg/mL. The GeoMx™ Nuclear Stain Morphology Kit (cat. 121300303, Bruker Spatial Biology, Seattle, WA) was used for nuclear staining with SYTO 13.

Following all staining steps, the slides were loaded onto the GeoMx™ DSP. All instrumentation steps were performed following steps outlined in the GeoMx™ DSP Instrument User Manual (MAN-10152). The GeoMx™ DSP was loaded with Instrument Buffer Kit (cat. 100474, Bruker Spatial Biology, Seattle, WA). Once slides were loaded and processed, a high-definition fluorescent image of the tissue was displayed by the instrument. Regions of interest (ROIs) were identified for collection. Each ROI was collected in at least triplicate and had over 200 nuclei for each replicate. Exceptions were made for nuclei counts that were lower, but in areas that were integral points of investigation. ROIs were further segmented into discrete areas of illumination (AOIs) to identify specific cell types using morphology marker masking. Three AOIs were defined for this project (PanCK+ cells, vimentin+ cells, remaining DNA signal).

After ROI/AOI selection, targeted UV light was directed by the GeoMx™ at each AOI releasing the UV-photocleavable oligonucleotide probes and via a microcapillary tube, probes from each AOI were dispensed into unique wells within a 96-well collection plate (cat. 100473, Bruker Spatial Biology, Seattle, WA).

Collected probes were taken through PCR and library prep according to GeoMx™ DSP NGS Readout User Manual (MAN-10153) to create a hybridized product of cleaved oligonucleotide probe aspirates with GeoMx™ SeqCode primers (Plates D, E, and F, Cat. 121400205 and 121400206, Bruker Spatial Biology, Seattle, WA) compatible with Illumina and Element AVITI platforms for Next Generation Sequencing. The GeoMx™ SeqCode primers are well-specific and include Illumina i5 and i7 dual indices which allow for spatial resolution by pairing an AOI with a specific well number, tying sequencing information back to the appropriate collection area. AMPure XP Beads (cat. A63880 Beckman Coulter, Brea, CA) were used for library purification. Quality checks on the pooled and purified library were carried out via qPCR on Light Cycler 480-II (Roche, Indianapolis, IN) using a KAPA Library Quantification (cat. KK4854, Roche, Indianapolis, IN) to measure concentration and the Agilent 5200 Fragment Analyzer System (Agilent Technologies, Santa Clara, CA) for library sizing using the NGS Fragment Kit (1-6000bp) (cat. DNF-473-0500, Agilent, Technologies, Santa Clara, CA). Samples were loaded in equimolar concentration and sequenced on the AVITI Sequencer (part #05 015 278 001, Element Biosciences, San Diego, CA) using a read configuration of 2x28bp with dual 8bp indexing, with a 2x75 Cloudbreak Freestyle Medium Sequencing kit (part #860-00014, Element Biosciences, San Diego, CA). FASTQ files generated from the AVITI were processed through the GeoMx™ NGS Pipeline, a Bruker Spatial software, that converts FASTQ files to digital count conversion (DCC) files. These converted files contain the sequencing information mapped to each specific well or AOI on the slide for in-depth spatial analysis using the built-in analysis software on the instrument or other analysis pipelines.

#### Raw data processing

Raw DCC files were imported with the canine whole transcriptome probe configuration file (Cf_R_NGS_CA_v1.0.pkc) and the annotation worksheet using readNanoStringGeoMxSet from GeomxTools. Counts of zero were shifted to one prior to quality control. Segment-level quality control applied the following thresholds: minimum 1,000 raw reads, at least 80% of reads trimmed, at least 80% stitched, at least 75% aligned, at least 50% sequencing saturation, minimum negative control count of 1, minimum 20 estimated nuclei, and minimum segment area of 1,000 square micrometers. Segments failing any evaluable criterion were removed. Probe-level quality control removed probes with a probe ratio below 0.1 or flagged as global Grubbs outliers, with local outlier removal enabled at a 20% failure threshold. Probes were collapsed to gene targets by geometric mean, yielding 1,963 targets. A per-segment limit of quantitation (LOQ) was defined as the negative probe geometric mean multiplied by the geometric standard deviation squared, with a floor of 2 counts. Segments detecting fewer than 10% of targets above the LOQ were removed, as were targets detected in fewer than 10% of segments, leaving 1,528 targets. Counts were normalized to the 75th percentile (Q3) of each segment; background normalization to the negative probe geometric mean was computed in parallel for comparison. *<u>Data processing</u>*. Segments were transferred to standR for batch correction^16^. Per-ROI quality control and a nuclei threshold of 20 were applied, and counts were TMM normalized. Principal component analysis showed clear separation by slide, so the 300 genes least variable with respect to slide were selected as negative control genes with findNCGs and RUV4 correction was applied with geomxBatchCorrection using AOI segment type as the biological factor. ROIs were placed on regions annotated as normal bladder, glandular, microcystic-tubular, or sarcomatoid. *<u>Differential expression</u>*. Differential expression used the limma-voom pipeline described in Section 1. The design was a means model on the combination of case and AOI segment (∼ 0 + biology) with RUV4 factors of unwanted variation as covariates. Contrasts compared the sarcomatoid case to the microcystic/conventional-urothelial case within the PanCK, vimentin, and CD45 compartments separately. A parallel contrast collapsing across AOI segment (∼ 0 + patient) provided a pseudo-bulk case-level comparison.

### Human bulk expression array

#### Data processing

For the MD Anderson SUC cohort^17^, Raw IDAT files for 112 samples were downloaded from Gene Expression Omnibus accession GSE128192 and read with readIDAT from the illuminaio package. Bead-type mean intensities were extracted per bead address, log2 transformed after adding one, and quantile normalized across arrays with normalizeBetweenArrays. Bead addresses were mapped to gene symbols using the GPL14951 platform annotation; probes without a symbol were discarded, and where multiple probes mapped to the same symbol the probe with the highest mean intensity across samples was retained. Sample identifiers were mapped to the descriptive sample names distributed with the series. Sarcomatoid and conventional urothelial carcinoma labels were taken from those sample annotations. Basal or double-negative molecular subsets were not included in the public deposit.

### Mouse BBN cell line RNA-seq

#### Dataset processing

Four count matrices were analyzed: three in vitro experiments profiling the BBN964, BBN966, and BBN975 lines before and after recycling with four replicates per condition, and one in vivo experiment profiling tumors from all three lines. Counts were scaled to reads per million by dividing each column by its library size in millions. Genes with zero variance across samples were removed. Gene-wise z-scores were computed by centering and scaling each gene across samples, and were used for visualization only.

### Signature based analyses

#### Molecular subtype classification

Consensus molecular subtypes of muscle-invasive bladder cancer were assigned with the consensusMIBC package^18^ using HGNC symbols, applied to normalized expression from each dataset. *<u>Pathway and transcription factor activity.</u>* Sample-level Hallmark pathway activity was inferred with VIPER^19^. Each MSigDB Hallmark gene set^20^ was converted to a regulon in which all member genes were assigned a mode of regulation of +1 and a likelihood of 1, and activity was computed with scale-based normalization. Transcription factor activity was inferred by the same approach using DoRothEA regulons restricted to confidence levels A, B, and C. Human regulons were used for all species. Features were ranked by variance across samples for visualization. *<u>Gene set enrichment for GeoMx.</u>* Hallmark gene sets were retrieved from MSigDB and gene sets with five or fewer genes represented on the panel were discarded^20^. Enrichment was tested with fry, a rotation-based test that accounts for inter-gene correlation, using the same design, contrast, and voom weights as the corresponding differential expression analysis. *<u>Immune deconvolution.</u>* Immune composition was estimated with CIBERSORT using the LM22 leukocyte signature matrix^21^. Normalized expression matrices were supplied as the mixture file and quantile normalized prior to support vector regression. The human LM22 signature was applied unchanged to canine and mouse data after symbol harmonization. Spatial deconvolution of the GeoMx data was additionally performed with SpatialDecon using a bladder cell profile matrix derived from GSE145137 and per-segment nuclei counts.

### Statistical details

Comparisons of individual genes, Hallmark activity scores, and deconvolved cell fractions between sarcomatoid and conventional groups were made by Welch two-sample t-test, with samples assigned to groups by their annotation label. Comparisons across the three in vivo BBN lines were made by one-way ANOVA. These tests were performed independently per feature and are reported as unadjusted p-values. Group comparisons of individual genes, pathway scores, and cell fractions were made by Welch two-sample t-test; comparisons across the three in vivo lines by one-way ANOVA.

### Anti-PD1 experiments

Mice were 10 to 12 weeks old C57BL/6J female mice (Strain 000664, The Jackson Laboratory, Bar Harbor, ME, USA) at time of initial injection. We housed animals for 2 weeks or more in our animal room after purchase. We subcutaneously injected 5×10^6 BBN966 or BBN964 cells (BBN966; Day -7, BBN964; Day -20). After tumors could be palpable (BBN966; 7 days later, BBN964; 20 days later), we randomly separated mice into two groups to achieve balanced tumor volumes, with each group containing 10 mice. On the same day, we treated mice with either anti-moue PD1 antibody (Cat#BE0146, Clone RMP1-14, Lot# 949225J3, Bio X cell, Lebanon, NH, USA) or Rat IgG2a isotype control (Cat#BE0089, Clone 2A3, Lot# 951625J2, Bio X cell) by intraperitoneal injection (Day 0). Treatment amount was 200ug in 100uL per injection. We treated and measured tumor size every three days until harvest day (BBN966; Day15, BBN964; Day26). In the BBN964 experiment, one mouse in the isotype control group, and two mice in the anti-mouse PD1 antibody group, died of unknown causes during the experimental period.

## Results

### Adaptive immunity restricts growth of basal UC tumors

In order to study the tumor microenvironment (TME) and create models to test the effectiveness of immunomodulatory agents in preclinical models of UC, we implanted basal UC tumor cell lines (5x10^6^, s/q), generated from treating immune competent (C57BL/6, e.g. B6) mice with oral *N-*Butyl-*N*-(4-hydroxybutyl)-nitrosamine (BBN)^22^, into naïve syngeneic B6 mice. This resulted in either complete lack of growth (BBN964, BBN966) or slow growth (BBN975) (**Fig. 1, left**). To increase tumorigenicity, we implanted tumor cells from these three cell lines into Rag1-KO mice as described^23^. As expected, in each case, tumors grew robustly in Rag1-KO mice (**Fig. 1, right**).

To determine whether tumor propagation in Rag1-KO tumors conferred tumor growth capacity in B6 mice, we passaged single cell suspensions from tumors grown in Rag1-KO mice directly into B6 mice (Rag1-KO -> B6). All three cell lines passaged in Rag1-KO mice had at least some tumor growth upon transfer to B6 mice, with BBN966 having the most robust growth **(Fig. 1, right)**. After repeated passaging in B6 mice, implanted BBN966 and BBN964 tumors had virtually 100% engraftment rate, with BBN966 notably more aggressive than BBN964 (**Fig. 9C, E**). We found that both BBN966 and BBN964 tumor cells plated on plastic after harvest subsequently grew tumors upon implantation (**Sup. Fig. 1A-B**). Finally, we found that BBN975 tumors, which grew upon transfer from Rag1-KO to B6 mice, did not grow upon subsequent transfer to a second B6 mouse (Rag1-KO -> B6 -> B6). Thus, passaging BBN cell lines in Rag1-KO mice augmented tumorigenicity which facilitated growth in immunocompetent mice.

### BBN derived tumors acquire variant histology upon passaging in immune deficient mice

To further characterize the recycled BBN964, BBN966, and BBN975 tumors, we asked three board certified pathologists with expertise in genitourinary oncology to blindly evaluate digital photographs of tumor sections stained with hematoxylin and eosin (**Fig. 2A, Sup. Fig. 2**). Tumors implanted in Rag1-KO mice acquired variant subtype histology, as most BBN966 tumor sections were either sarcomatoid or sarcomatoid squamous, BBN975 tumor sections were either squamous or sarcomatoid squamous, and BBN964 were either sarcomatoid or squamous (**Fig. 2B, Sup. Table 1**). Similarly, tumors recycled in Rag1-KO for a second time were diagnosed as sarcomatoid (BBN966), squamous or sarcomatoid squamous (BBN975), or sarcomatoid or squamous (BBN964) (**Fig. 2C, Sup. Table 2**). This was consistent with tumors passaged in Rag1-KO then transferred to wild type mice (C57BL/6J, e.g. B6) (**Fig. 2D, Sup. Table 3**). Thus, BBN derived tumor cells passaged in mice lacking adaptive immunity acquired variant subtype histology, and tumors were generally classified as sarcomatoid (BBN966), squamous (BBN975), or a hybrid of the two (BBN964).

**Figure 2.**
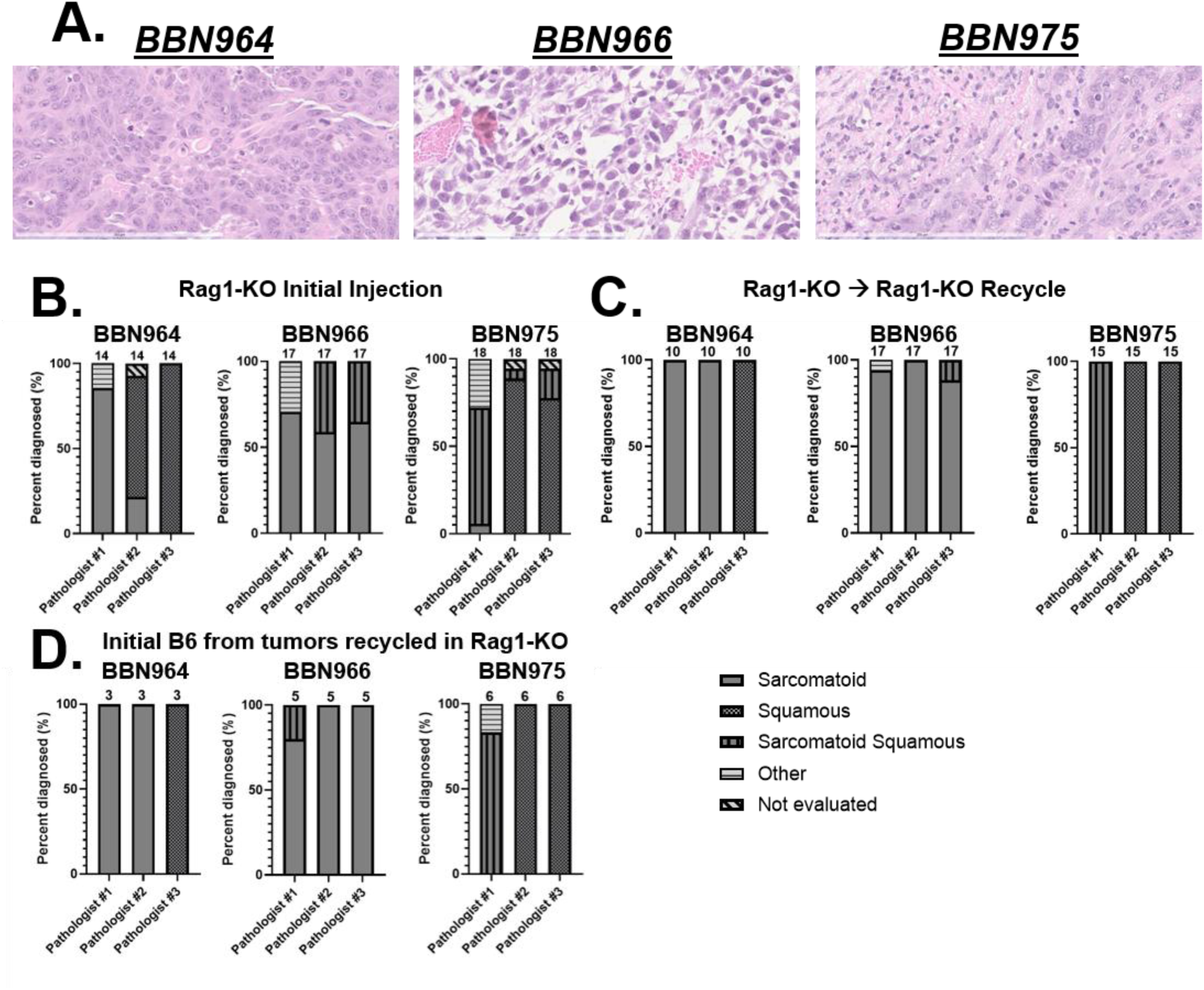
Passaging basal tumors in adaptive immune deficient mice results in tumors with variant subtype histology. **A.** Representative pictures of BBN964 (left), BBN966 (middle), and BBN975 tumors (right). **B-D.** Blinded pathologic diagnoses from three independent GU oncology pathologists from photographs of hematoxylin and eosin (H and E) slides generated from BBN964, BBN966, and BBN975 tumors. The number at the top of each bar represents how many diagnoses were made from unique pictures per pathologist. **B.** Pathologic diagnoses performed on tumors initially implanted (first injection) into Rag1-KO mice. **C.** Pathologic diagnosis performed on tumors recycled into Rag1-KO mice (second injection) after initial implantation into Rag1-KO mice. **D.** Pathologic diagnoses performed on tumors implanted in B6 following two passages in Rag-1 KO (BBN964 and BBN966), and tumors implanted in B6 following one passage in Rag1-KO (BBN975).

### SUC tumors in human, dog, and mouse express low Keratin 5

To classify the recycled BBN964, BBN966, and BBN975 tumors based on gene expression profiling, we performed immunofluorescence on bulk tumors passaged in B6 mice and RNAseq on tumor cells. Initially, we explored the role of the basal marker keratin 5 (*Krt5*), as this gene was highly expressed in basal UC and basal SUC, but not expressed in double negative SUC tumors, which were associated with low expression of basal and luminal markers, worse survival, and higher expression of EMT genes^17^. We found low to negative expression of Keratin 5 in bulk BBN966 tumors by immunofluorescence, which was significantly lower than Keratin 5 expression in bulk BBN964 and BBN975 tumors (**Fig. 3A**). While BBN964 and BBN975 tumor cells downregulated *Krt5* following passage in B6 mice, BBN966 tumor cells expressed near zero *Krt5* before and after passage in B6 mice that was far lower than post-recycling BBN964 and BBN975 values (**Fig. 3B**). To confirm low Keratin 5 as a characteristic of SUC, we compared expression of *KRT5* utilizing human RNAseq datasets. Utilizing a publicly available RNAseq dataset of human SUC and UC tumors^17^, we found that *KRT5* expression was significantly lower in SUC tumors (**Fig. 3C**). Next, we assessed *KRT5* expression in our cohort, which entailed spatial transcriptomic profiling of SUC and UC tumor regions of interest (ROI) in tumors with mixed SUC and UC histology, in order to compare SUC and UC ROI within the same tumor^15^. Similarly, we found that SUC ROI had significantly lower *KRT5* expression than UC regions (**Fig. 3D**). To confirm low Keratin 5 expression was a cross-species phenomenon in SUC, we assessed spatial transcriptomics data utilizing multiple ROI from one dog with SUC and UC in the same tumor. We found that SUC ROI had significantly lower *Krt5* expression when compared with UC ROI (**Fig. 3E**). Thus, low Keratin 5 expression was found in SUC across species, including human, dog, and BBN966 SUC tumors.

**Figure 3.**
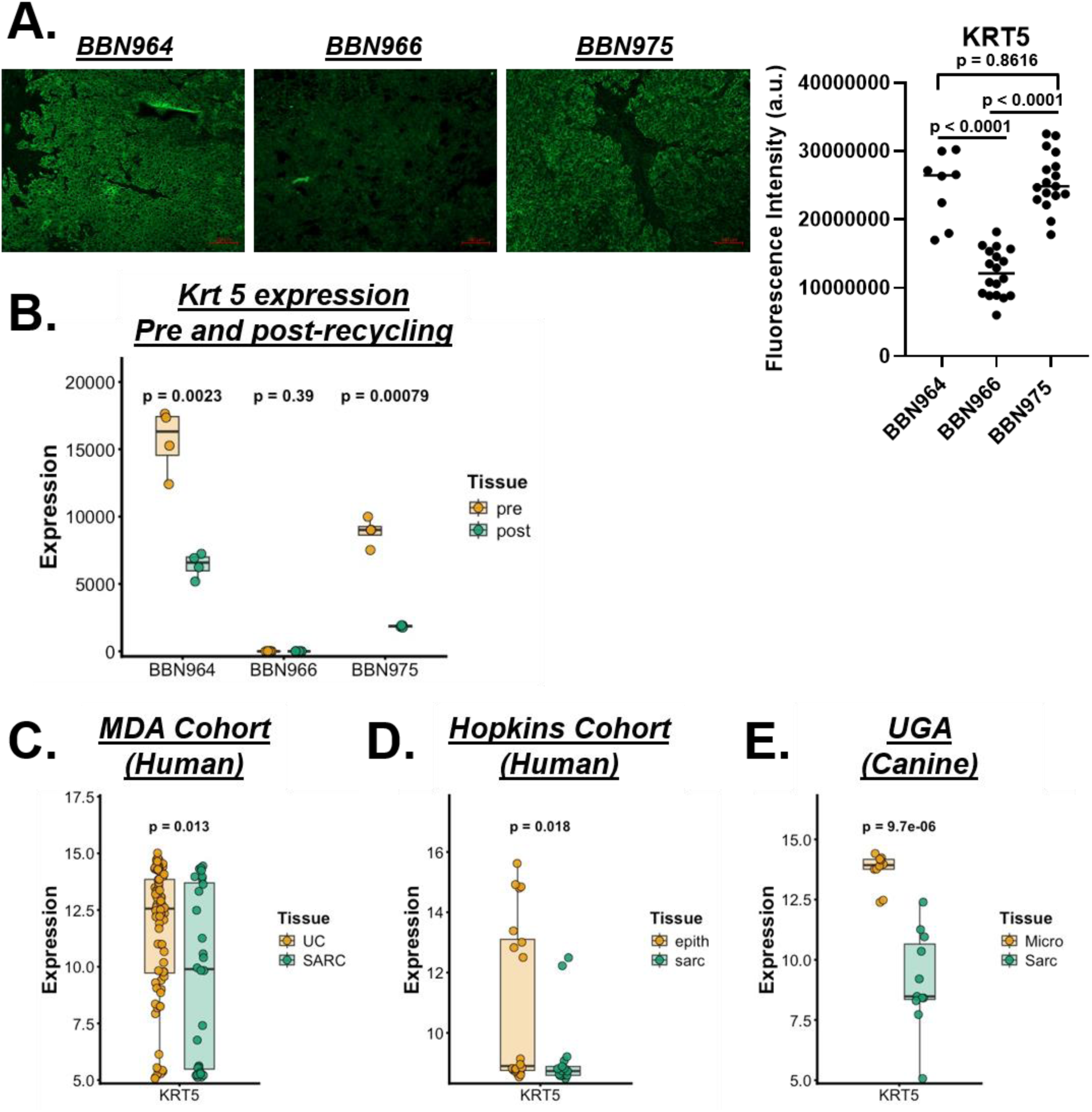
Low keratin 5 expression is a cross-species hallmark of SUC. **A.** BBN964 (hybrid), BBN966 (SUC), and BBN975 (squamous) tumors were stained via immunofluorescence for Keratin 5. Left, Representative photos taken from pictures with the median fluorescence intensity within each tumor group. Right, plot of the fluorescence intensity of all photos taken from slides stained for Keratin 5. **B.** Expression of *Krt5* on parental (pre-) and recycled (post-) tumor cells for BBN964, BBN966, and BBN975 obtained from RNA sequencing. **C.** *KRT5* expression in the MD Anderson cohort of separate sarcomatoid and conventional urothelial carcinoma tumors, via RNA sequencing. **D.** *KRT5* expression in the Hopkins cohort of sarcomatoid and urothelial carcinoma tumors within the same patients, from spatial transcriptomics data of unique regions of interest (ROI). **E.** *Krt5* expression from UGA spatial transcriptomics of different ROI from one canine patient with mixed UC and SUC.

### BBN966 tumors lack expression of basal and luminal markers

To explore whether recycled tumor cells match the genetic profile of human SUC cells, we interrogated BBN964, BBN966, and BBN975 tumor cells for luminal and basal UC markers. Human SUC tumors were found to be negative for luminal markers, and previously subdivided into groups based on positive and negative expression of basal markers (basal SUC and double negative SUC, respectively)^17^. Using the MD Anderson dataset, we replicated this result (**Sup. Fig. 3A**), and using unsupervised clustering, found that most human SUC and UC clustered separately (**Sup. Fig. 3B**). Similarly, in our dataset of SUC and UC ROI within the same human tumors, we found that most SUC and UC clustered separately by unsupervised hierarchical analysis (**Sup. Fig. 3C**). Upon separation of SUC and UC ROI, the SUC component appeared to have relatively low expression of luminal and basal markers (**Fig. 4A**). When comparing parental (pre) with recycled (post) tumor cells, many luminal and basal markers were downregulated in recycled cells across all three murine models (BBN964, BBN966, BBN975, **Fig. 4B-D**). However, while recycled BBN964 and BBN975 tumors exhibited upregulation of several luminal markers, recycled BBN966 tumors exhibited downregulation of almost all luminal (including *Gata3, Krt7, Krt8, Krt18, Krt19, Foxa1*) and basal (including *Cdh3, Krt14, Pparg, Cd44*) when compared with parental tumor cells (**Fig. 4B-D**).

**Figure 4.**
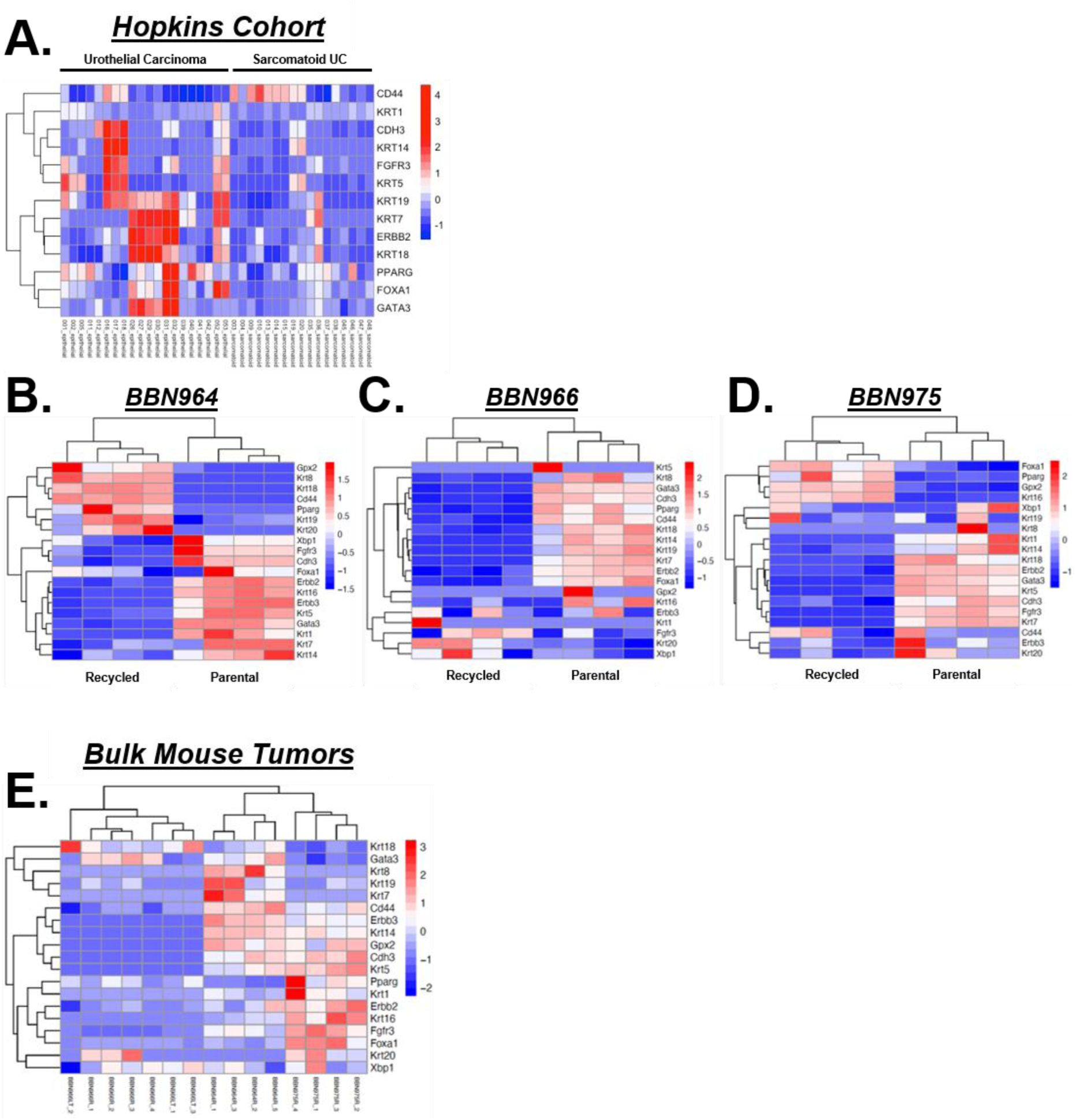
Consistent with a cohort of human SUC, mouse SUC tumors exhibit a “double-negative” phenotype. **A.** Supervised clustering of the Hopkins cohort of sarcomatoid and urothelial carcinoma tumors within the same patients, from spatial transcriptomics data of unique regions of interest (ROI). **B-D.** Unsupervised clustering of luminal and basal gene expression from RNAseq evaluation of parental (non-tumorigenic) and post-recycling (tumorigenic) tumor cells (BBN964, left; BBN966, center; BBN975, right). Recycled is on the left, parental is on the right. **E.** Unsupervised clustering of luminal and basal genes from RNAseq of bulk mouse tumors from BBN964, BBN966, BBN966 passaged in B6 mice for ∼1 year (BBN966LT), and BBN975.

To determine whether BBN964, BBN966, and BBN975 tumors were genetically distinct, we performed unsupervised clustering on RNAseq from bulk tumors from B6 mice, and assessed for luminal and basal marker expression. We found that BBN964, BBN966, and BBN975 tumors clustered separately (**Fig. 4E**). BBN966 tumors from initial passage in B6 mice (BBN966R) clustered with BBN966 tumors passaged in B6 mice for approximately 1 year (BBN966LT), suggesting that BBN966 tumors maintained their phenotype long-term (**Fig. 4E**). BBN966 tumors generally had lower expression of luminal and basal markers than BBN964 and BBN975 tumors (**Fig. 4E**). Thus, BBN964, BBN966, and BBN975 were genetically distinct tumors, consistent with their unique growth kinetics (**Fig. 1 and 9C-D**) and phenotype by genitourinary pathologist assessment (**Fig. 2**).

### High expression of genes upregulated in recycled tumors correlates with worse OS

Next, we further explored changes in the tumor cells before recycling (parental) and after acquiring growth capacity in B6 mice (recycled). Using principal components analysis, we found that all three tumor cells (BBN964, BBN966, BBN975) were genetically distinct from one another, and all three cell lines underwent distinct genetic changes upon recycling such that these cells were distinct from the parental cell line in each case (**Fig. 5A**). There were many genes upregulated and downregulated when comparing parental with recycled cells (**Fig. 5B**), including genes upregulated (**Fig. 5C, top**) and downregulated in recycled cells (**Fig. 5C, bottom**) that are unique to each cell line, along with genes that were upregulated (328) and downregulated (106) in all three cell lines. Bootstrapping analysis demonstrated that these changes did not occur by chance (**Fig. 5D**).

**Figure 5.**
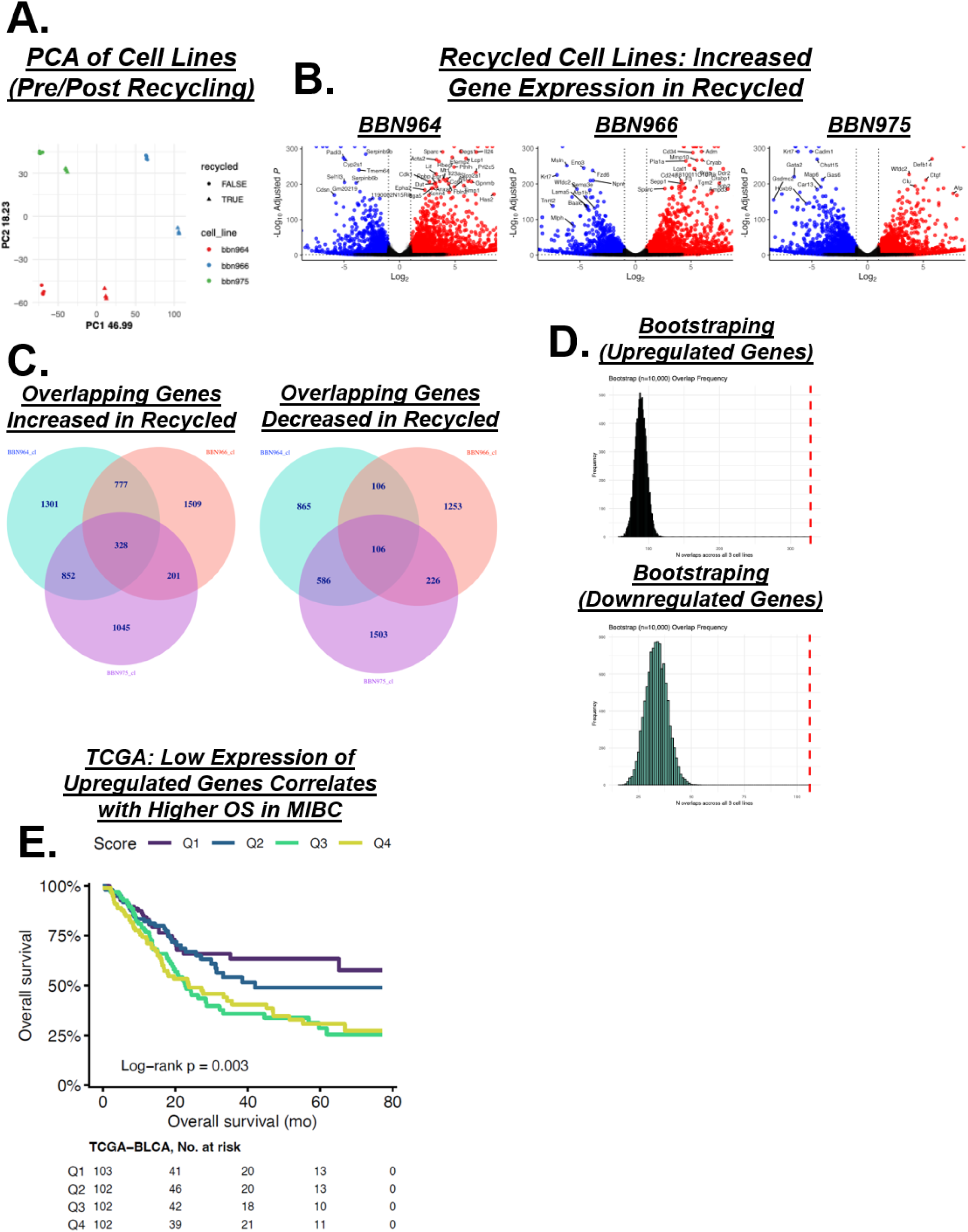
High tumor expression of genes upregulated in all three mouse tumor lines correlates with worse overall survival in patients with muscle invasive UC. **A.** Principal components analysis of BBN964 (red), BBN966 (blue), and BBN975 (green) RNA sequencing analysis of parental (circle) and recycled (triangle) cells. B. Volcano plots of BBN964 (left), BBN966 (center), and BBN975 (right) comparison of upregulated genes (red) and downregulated genes (blue) when comparing parental versus recycled tumor cells. **C.** Venn diagram of genes upregulated (left) and downregulated (right) in recycled tumor cells. There were 328 unique genes upregulated in all 3 recycled tumor cell groups, and 106 unique genes downregulated in all 3 recycled tumor cell groups. **D.** Bootstrapping analysis demonstrated number of genes expected to be upregulated (top) and downregulated (bottom) in all 3 recycled tumor cell groups if the number of genes was due to chance. Red line denotes number of genes observed to be upregulated in all 3 recycled tumor cell groups (328, top) and number of genes observed to be downregulated in all 3 recycled tumor cell groups (106, bottom). **E.** Tumors from patients with muscle invasive urothelial carcinoma from The Cancer Genome Atlas (TCGA) were divided into quartiles based on lowest (1) to highest (4) expression of genes upregulated in all 3 tumor cell lines, then overall survival graphed in a Kaplan-Meier curve.

To determine whether information obtained from the acquisition of tumorigenicity in the BBN964, BBN966, and BBN975 tumor cells has clinical relevance, we created a scoring system utilizing the 328 genes upregulated in all three recycled cell lines. Next, we segregated muscle invasive bladder cancer tumors from The Cancer Genome Atlas (TCGA) by expression of these genes. We found that patients who had tumors with the lowest quartile of expression of genes upregulated by recycled cells had significantly higher overall survival (OS), while patients with the highest gene expression had lower OS (**Fig. 5E**). Thus, identification of genes associated with tumorigenicity in mouse BBN derived tumors can be utilized to identify patients with less aggressive bladder cancer and greater OS.

### Increased EMT is a hallmark of human SUC and all recycled tumors

To identify pathways upregulated in recycled mouse tumor cells, we assessed recycled tumor cells from B6 mice by GSEA pathway analysis. We noted the EMT transition pathway was upregulated in all 3 recycled tumor cells, including sarcomatoid (BBN966), squamous (BBN975), and hybrid (BBN964) (**Fig. 6A**). To further characterize EMT in recycled mouse tumor cells, we plotted the hallmark EMT score^20^ for BBN964, BBN966, and BBN975, such that a higher EMT score was associated with a more mesenchymal tumor. As noted above, and consistent with a more advanced EMT state, all recycled tumor cells had higher EMT score when compared with parental tumor cells (**Fig. 6B**).

**Figure 6.**
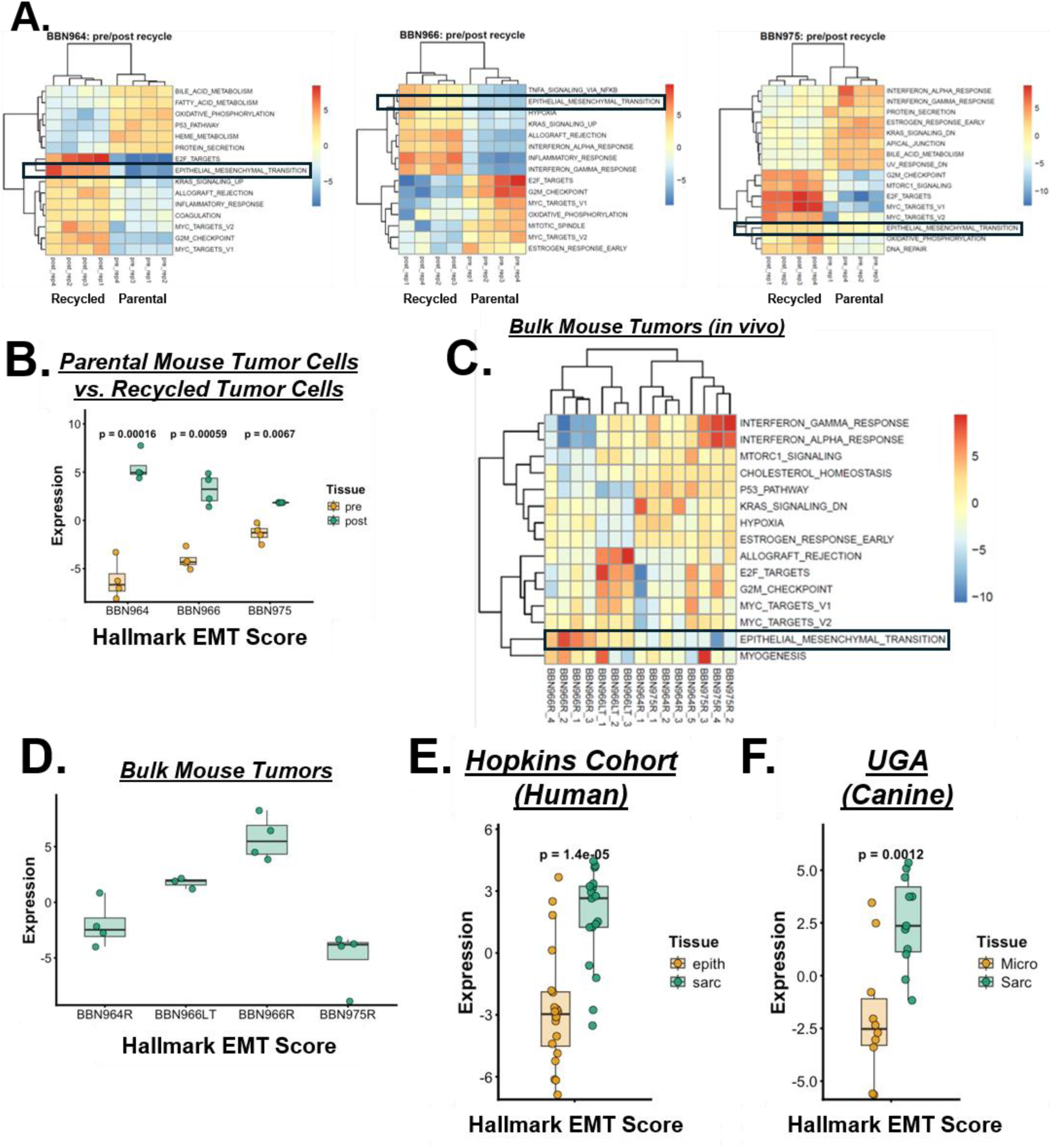
Epithelial to mesenchymal transition is a cross-species hallmark of SUC. **A.** Pathway analysis generated via analysis of RNA sequencing performed on parental (right) and recycled (left) tumor cells from BBN964 (left), BBN966 (center), and BBN975 (right) tumor cells. **B.** Hallmark EMT score for parental and recycled tumor cells obtained by RNA sequencing. **C.** Unsupervised clustering of pathway analysis of bulk *in vivo* generated BBN964, BBN966, BBN966 tumors passaged in mice for ∼1 year (BBN966LT), and BBN975 tumors. **D.** Hallmark EMT score for bulk tumors obtained by RNA sequencing. **E-F.** Hallmark EMT score for **(E.)** the Hopkins cohort of urothelial carcinoma (epith) and sarcomatoid urothelial carcinoma (sarc) from multiple ROI of mixed tumors, obtained via spatial transcriptomics, and **(F.)** the UGA canine tumor of mixed urothelial carcinoma (Micro), and sarcomatoid urothelial carcinoma (Sarc), obtained via spatial transcriptomics of multiple ROI.

Next, we explored upregulated pathways in bulk SUC tumors, and found both the BBN966 and BBN966LT tumors had the highest EMT pathway expression when compared with other groups (**Fig. 6C**). This was consistent with BBN966 having the highest hallmark EMT score (**Fig. 6D**). As EMT has been described as a hallmark of human SUC^17^, we also examined upregulated pathways in human SUC, using the Hopkins cohort of patients with mixed UC and SUC initially described in **Fig. 3D**. We found the SUC ROI had increased EMT pathway expression (**Fig. 6E, Sup. Fig. 4A**). Finally, we explored pathway analysis from spatial transcriptomics data from canine tumors with mixed UC and SUC (described in **Fig. 3E**) and found that EMT was consistently upregulated in SUC (**Sup. Fig. 4B**), with a hallmark EMT score that was significantly higher in canine SUC than UC (**Fig. 6F**). Thus, we found increased EMT in all recycled mouse tumor cells, as well as increased EMT in canine and human SUC.

### EMT regulators p63 and E-cadherin have low expression in murine SUC

TP63 regulates expression of keratins such as *KRT5*^24^, and inhibits EMT progression^25^. As we found *Krt5* was downregulated (**Fig. 3**) and EMT associated genes were upregulated (**Fig. 6**) in BBN966, we explored the role of p63 expression in our recycled mouse tumors by staining BBN964, BBN966, and BBN975 bulk tumors for p63. BBN966 bulk tumors had significantly less expression of p63 compared with BBN964 and BBN975, with BBN964 having intermediate expression between BBN966 (lowest) and BBN975 (highest) (**Fig. 7A**). Consistent with this, BBN966 has low expression of *Tp63* in both parental and recycled tumor cells that was lower than BBN964 and BBN975 (**Fig. 7B**).

**Figure 7.**
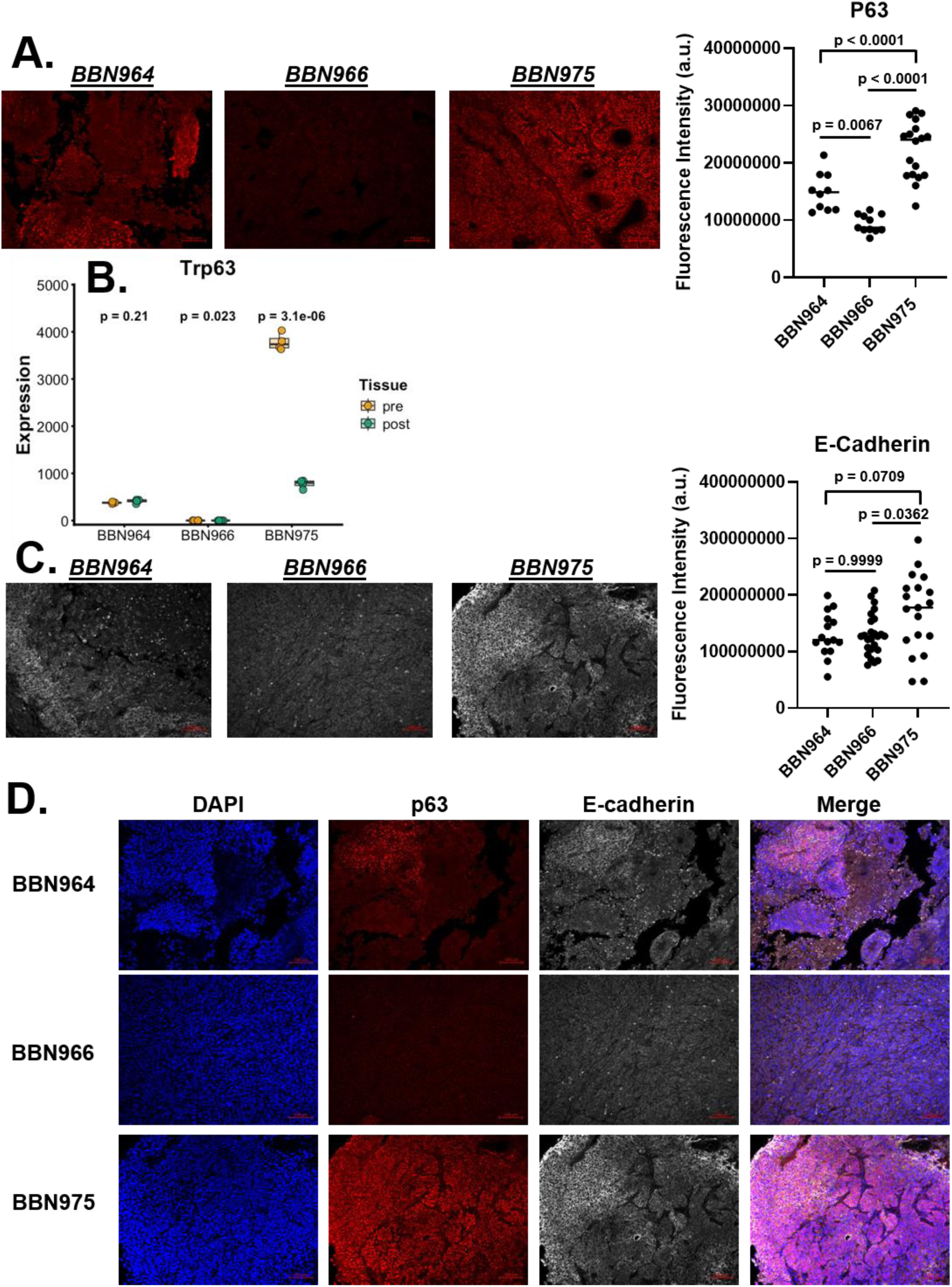
Low p63 and E-cadherin expression in mouse SUC. **A.** BBN964 (hybrid), BBN966 (SUC), and BBN975 (squamous) tumors were stained via immunofluorescence for p63. Left, Representative photos taken from pictures with the median fluorescence intensity within each tumor group. Right, plot of the fluorescence intensity of all photos taken from slides stained for p63. **B.** Expression by RNA sequencing of *Tp63* in parental (pre-) and recycled (post-) tumor cells for BBN964, BBN966, and BBN975. **C.** BBN964 (hybrid), BBN966 (SUC), and BBN975 (squamous) tumors were stained via immunofluorescence for E-cadherin. Left, Representative photos taken from pictures with the median fluorescence intensity within each tumor group. Right, plot of the fluorescence intensity of all photos taken from slides stained for E-cadherin. **D.** For BBN964 (top), BBN966 (middle), and BBN975 (bottom), representative photos of DAPI (nuclear stain, blue), p63 (red), E-Cadherin (white), and Merged overlay images. Note for BBN975 the tumor photograph used is the same as for **C.** as the goal was to show tumors with fluorescence values as close to the median for p63 and E-Cadherin as possible. P values for p63 and E-cadherin graphs were calculated using Tukey’s multiple comparisons test.

As E-cadherin downregulation is also associated with EMT, we also examined E-cadherin expression in bulk recycled mouse tumors, and we found BBN966 had relatively low expression of E-cadherin, comparable to BBN964, and significantly lower than BBN975 (**Fig. 7C**). Finally, we examined p63 and E-cadherin expression within the same tumor slide, which again demonstrated low p63 and E-cadherin expression in BBN966 when compared with other tumors (**Fig. 7D**).

### Human and mouse SUC are enriched for M2 macrophages

Human SUC appeared to have an abundance of macrophages with suppressive M2 phenotype per CIBERSORT, when compared with pure UC, though these were not formally compared between groups^17^. Thus, we compared abundance of macrophage subtypes between human SUC and UC, using CIBERSORT analysis. Human SUC tumors have significantly more macrophages with a M2 gene expression phenotype than UC tumors, and M2 macrophages in SUC outnumber M0 and M1 macrophages (**Fig. 8A**). Similarly, when comparing SUC and UC within the same tumor, there were significantly more M2 macrophages in SUC (**Fig. 8B**). Unlike SUC and UC from unique patients, SUC tumors had significantly more M0 macrophages when comparing SUC and UC within the same tumor (**Fig. 8B**). Thus, human SUC tumors were enriched for M2 macrophages.

**Figure 8.**
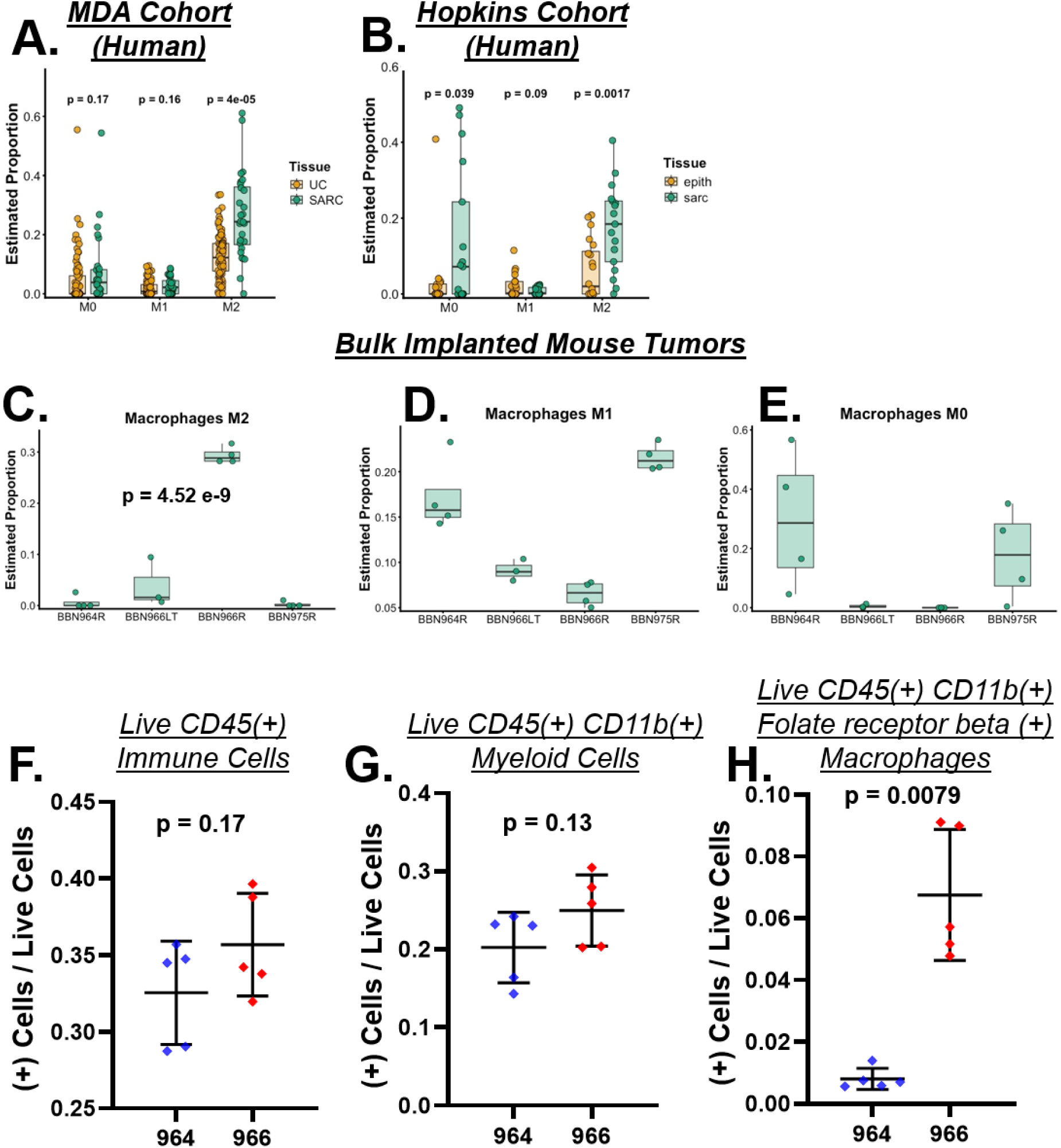
M2 macrophages are enriched in human and mouse SUC. **A.** M0, M1, and M2 macrophage gene expression in SUC and UC from the MD Anderson cohort. **B.** M0, M1, and M2 macrophage gene expression between SUC and UC ROI in the Hopkins cohort. **C-E. (C.)** M2 macrophage gene expression, **(D.)** M1 macrophage gene expression, and **(E.)** M0 macrophage gene expression from RNA sequencing data of bulk tumors from recycled BBN964, BBN966, BBN966LT, and BBN975 tumors. **F-H. (F.)** CD45+ leukocytes, **(G.)** CD45+CD11b+ myeloid cells, and **(H.)** CD45+CD11b+ cells expressing folate receptor beta, as a proportion of total live cells in BBN964 and BBN966 tumors (both passaged in B6 for at least 1 year).

Next, we compared macrophage infiltration of implanted BBN964, 966, and 975 tumors via CIBERSORT analysis of RNAseq data from bulk tumors. Consistent with human SUC, BBN966 tumors had significantly higher M2 macrophages than BBN964 and BBN975 **(Fig. 8C**), and BBN966 had higher M2 macrophage proportion than M0 or M1 (**Fig. 8C-E**). Conversely, BBN964 and BBN975 had higher M1 macrophage proportion than BBN966, and variable M0 (**Fig. 8D-E**). We assessed BBN966 bulk tumors that have been passaged in mice for more than 1 year, and found decreased M2 macrophage gene expression compared with BBN966 that was not recycled long-term (**Fig. 8C**).

To further evaluate macrophages from tumors that had been passaged long term in mice, we evaluated myeloid cell infiltration of BBN964 and BBN966 tumors by flow cytometry from tumors that had been passaged in B6 mice for at least one year. Both BBN964 and BBN966 tumors were immune infiltrated, with CD45+ leukocytes comprising approximately 30-40% of live cells (**Fig. 8F**). We then assessed for CD11b+ myeloid cells (**Fig. 8G**), and the presence of folate receptor beta (FRb) on CD11b+ myeloid cells (**Fig. 8H**), as FRb is a specific M2 macrophage marker^26^. Consistent with the CIBERSORT analysis of BBN966 bulk tumors, we found FRb+CD11b+CD45+ M2 macrophages comprised a larger percentage of live BBN966 bulk tumor when compared with BBN964 (**Fig. 8H**). There was not a significant difference in B cells or T cells as a percentage of live cells when comparing BBN964 and BBN966 bulk tumors (**Sup. Fig. 5**). Thus, the BBN966 TME was enriched for M2 macrophages, which was consistent with human SUC.

### Human SUC has high PD-L1, driven by TPS score, and consistent with ICI inhibition of mouse SUC growth

High PD-L1 correlated with response and survival following ICI treatment in multiple tumor subtypes ^27–30^. Previous reports have described high PD-L1 gene expression (*CD274*) in a cohort of patient SUC tumors^3,17^, and recently we described a cohort of 22 patients with SUC where most tumors expressed high PD-L1 combined positive score (CPS) by immunohistochemistry^31^. To extend these results, we added an additional 16 patients to our published cohort, for a total of 38 patients, and found that 81.5% (31 of 38) had PD-L1 CPS score greater than 10 (**Fig. 9A**), considered a cutoff for positive score in patients with bladder cancer, ref. ^32^). Moreover, 52.6% (20 of 38) had very high PD-L1 CPS score of 50 or greater, suggesting high PD-L1 CPS is a hallmark of a significant subset of SUC tumors. For tumors where immune cell score (ICS) and tumor proportion score (TPS) were available (n = 17), the tumor proportion score was significantly higher than the immune cell score, suggesting that in most cases, TPS score is the primary driver of the PD-L1 CPS score (**Fig. 9B**).

**Figure 9.**
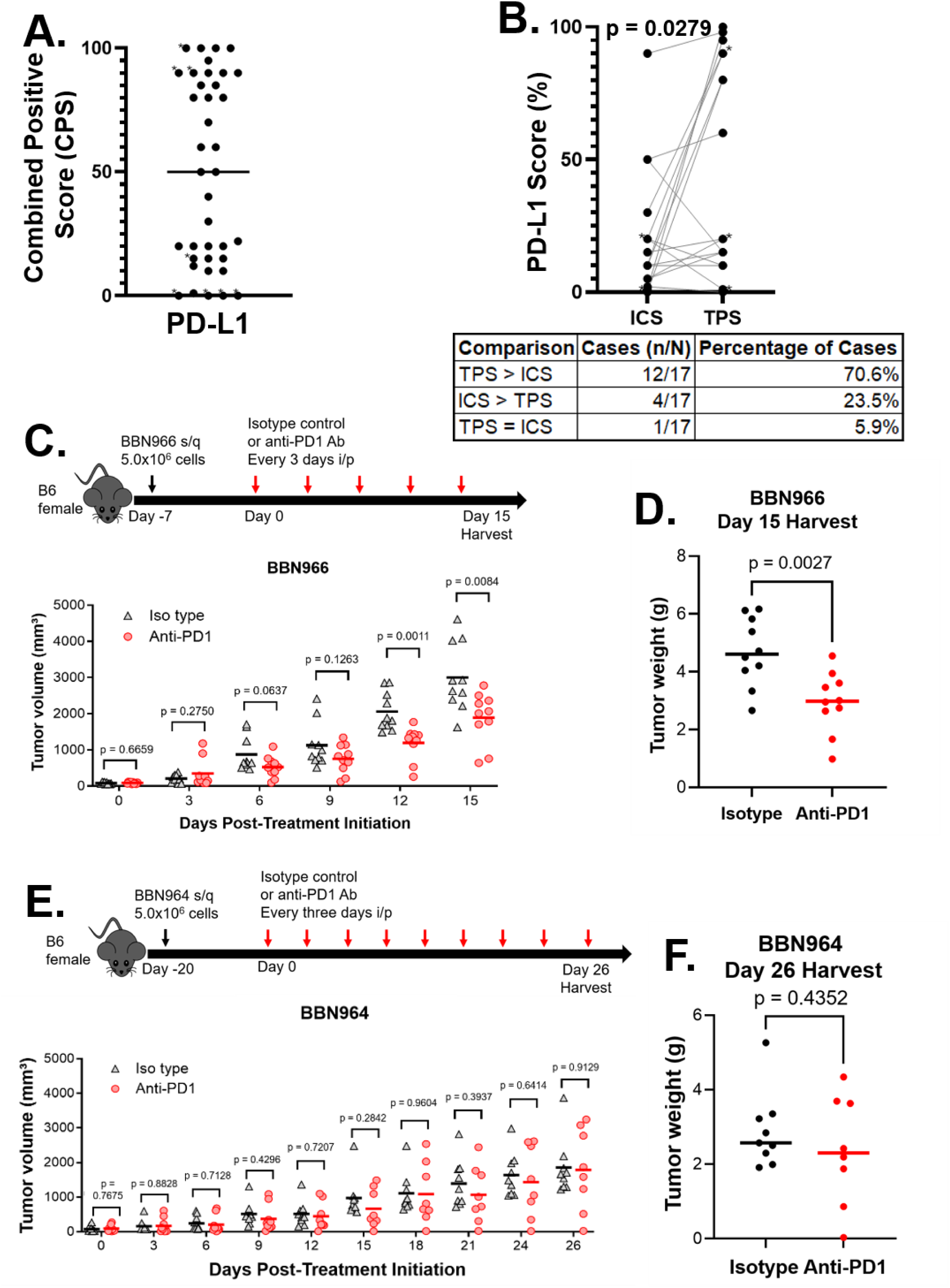
Mouse SUC is ICI responsive, and high PD-L1 score in human SUC is driven by tumor proportion score. **A.** PD-L1 CPS score on 38 patients with SUC. **B.** Immune cell score (ICS) and tumor proportion score (TPS) for patients in (A.) where both ICS and TPS were available (n = 17). TPS was significantly higher than ICS in SUC tumors (median difference = 10, Wilcoxon matched-pairs signed-rank test, n=17, P = 0.0279), with TPS exceeding ICS in 70.6% of cases (12/17). When values were reported as a range, we utilized the lower number as follows: ICS: 20-30 = 20 (2 patients), ICS: 1-5 = 1 (1 patient), TPS: 90-100 = 90 (1 patient), TPS: 20-30 = 20 (1 patient), TPS: <1 = 0 (2 patients), TPS: >90 = 90 (1 patient). **C-D.** BBN966 tumor experiment schema (**C.,** top). BBN966 tumors were implanted, then 7 days later (when tumors were approximately ∼100 mm^3) tumors were measured via calipers, separated into two groups (n = 10 per group) with mice with equivalent tumor volumes, then injected with either anti-PD1 or isotype control every 3 days. Tumors were measured every 3 days via calipers **(C.)**, harvested 15 days after starting treatment and weighed **(D.)**. **E-F.** BBN964 tumor experiment schema (**E.,** top). BBN964 tumors were implanted, then 20 days later (when tumors were approximately ∼100 mm^3) tumors were measured via calipers, separated into two groups (n = 10 per group) with mice with equivalent tumor volumes, and were treated as in **C-D.** above. Tumors were measured every 3 days via calipers **(E.)**, harvested 26 days after starting treatment, and weighed **(F.).** Note in the BBN964 experiment, one mouse from the isotype group and two mice from the anti-PD1 group died during treatment.

As others have reported a high response rate to ICI with SUC^4^, we asked whether anti-PD1 ICI impedes tumor growth in the BBN966LT SUC model. We implanted recycled BBN966 tumor subcutaneously into syngeneic (B6) mice, and when tumors were approximately 100 mm^3, we separated mice into equal groups and treated with intraperitoneal anti-PD1 antibody or isotype control every 3 days. We found that BBN966 tumors treated with anti-PD1 had significantly slower tumor growth (**Fig. 9C**) and smaller tumor weights (**Fig. 9D**) when compared with tumors from mice treated with isotype control. Next, to determine if that was a general phenomenon across all BBN UC derived tumors, we asked whether anti-PD1 antibody slowed tumor growth in BBN964LT tumors using a similar approach. Consistent with BBN964 being a less aggressive hybrid variant subtype histology, BBN964 tumors grew more slowly than BBN966 tumors (**Fig. 9E–F**). We found that BBN964 tumors had a variable response to anti-PD1, with a subset of tumor volumes smaller than control, and a subset of tumor volumes growing at similar rate as control (**Fig. 9E)**. We found that tumor weights mirrored the results obtained with caliper measurements (**Fig. 9F**). Thus, the BBN966 SUC tumors exhibits similar response to ICI as found in human SUC, while BBN964 had a mixed response. This was consistent with the different histologic diagnosis of BBN964 that was unique from BBN966 (**Fig. 2**).

## Discussion

SUC is a rare variant histologic subtype^33^ that is associated with worse OS^5,6^. Given limited treatment options, there is a need for preclinical models to test candidate treatment strategies. This report describeed a novel tumor cell line that matches human SUC in histologic appearance by expert GU pathologic diagnosis, via lack of expression of basal and luminal markers, high EMT pathway expression, high expression of genes consistent with M2 macrophage predominance, and response to anti-PD1 ICI. This preclinical model is complementary to the development of patient derived SUC organoids^13^, as while SUC organoids are superior to mouse models in modeling human disease, the lack of a complete immune system makes study of immunotherapy combinations difficult. As others have demonstrated that anti-PD1 ICI is active in a subset of SUC^4^, PD-L1 is high in a significant cohort of human SUC (**Fig. 9** and ^31^), and we demonstrated that anti-PD1 ICI inhibited tumor growth in BBN966 SUC (**Fig. 9**), future preclinical therapeutic study in SUC will likely necessitate testing of combination therapy with ICI and other immunomodulatory agents.

Our previous work suggested that macrophage gene expression was higher in human SUC than UC^15^. This work extended our previous work by demonstrating higher M2 macrophage gene expression in human SUC when compared with UC, and higher M2 macrophage enrichment in human SUC when compared with naïve (M1) or immunostimulatory (M1) macrophages. Our spatial transcriptomics data comparing mixed UC and SUC suggested transforming growth factor beta (TGFb) expression is higher in SUC^15^, which is consistent with a previous report^17^. As TGFb induces M2 macrophage polarization^34^, this may be how M2 macrophage enrichment occurs in SUC. TGFb signaling also regulates EMT and bladder cancer tumor invasiveness^35^. Thus, TGFb may facilitate M2 macrophage polarization and EMT in SUC. Although additional mechanistic work is needed to further clarify the functional nature and subtype of SUC macrophages^36,37^, recent work identified a SPP1+ macrophage subset as a critical immune regulator in bladder cancer^38^, suggesting the importance of these immune cells in regulating UC.

The identification of predictive and prognostic biomarkers in UC is an area of active study. We identified genes upregulated in all three mouse models when comparing parental tumor cells that are not tumorigenic, with recycled cells that are tumorigenic. Our finding that the patients with tumors with lowest expression of these genes had the greatest OS in the TCGA muscle invasive UC cohort is consistent with the notion that these genes, which are associated with tumorigenicity in our mouse models, also drive tumor aggressiveness in humans. It also provides further evidence that these mouse models are relevant to human disease, as high expression of these genes, which ostensibly drove acquisition of tumor growth capacity in mice, associated with decreased OS in human UC. Further directions include determining whether expression of these genes constitutes a program that associates or drives the presence of circulating tumor DNA in UC^39^, which is associated with decreased OS in high risk UC post-cystectomy but also identifies a subset of patients that may benefit most from ICI^40^, Also, on-treatment ctDNA clearance associated with higher OS in adjuvant muscle invasive UC^41^ and complete response in metastatic UC^42^. How tumor expression of these genes is associates with immune based biomarkers in UC and SUC is another area that could improve our mechanistic understanding of how tumorigenicity relates to immune infiltration^43^.

We noted high expression of EMT gene expression in recycled murine variant histology tumors, including SUC, which was consistent with acquired tumorigenicity noted in all recycled tumors (**Fig. 1**). Bulk tumors from the BBN966 model had the highest EMT, consistent with its aggressive growth (**Fig. 1 and 9C-D**). Low p63 expression in BBN966 SUC is consistent with prior reports demonstrating p63 inhibits EMT in bladder cancer^25^ and activates keratin 5 expression^24^. Low E-cadherin also drives tumor aggressiveness and is associated with EMT^44^, and E-cadherin activation inhibits tumor metastatic potential^45^. The lack of basal and luminal markers in BBN966 SUC, and a subset of the Hopkins cohort and MD Anderson SUC tumors, is consistent with attributes of a previously described claudin-low UC subtype distinct from conventional basal and luminal UC tumors, such as high EMT and high immunosuppression, though claudin-low tumors were classified as basal using conventional classifiers^46^. Also consistent with upregulated EMT in SUC, peroxisome proliferator activated receptor – gamma (Pparg), which inhibits EMT^47^, is downregulated in BBN966 SUC tumor cells post-recycling, as well as most SUC tumors in the Hopkins and MD Anderson cohorts. High EMT in tumors correlates with expression of high immune checkpoint proteins and immune suppression^48^, however the mechanism by which this occurs in SUC is poorly understood.

We demonstrated for the first time that high PD-L1 CPS score in SUC is driven primarily by the TPS. Previous work has shown that PD-L1 expressed by the tumor cell itself can block T cell function and facilitate immune suppression^49^. Thus PD-L1 on the tumor cell may facilitate immune suppression and tumor growth in SUC. The percentage of patients with SUC and PD-L1 CPS score ≥ 10 is higher (81.5%) than historical tumor data from patients with advanced UC (43%, ref. ^50^). Although PD-L1 is an inconsistent biomarker for ICI response in solid tumors, several FDA approved indications for the use of single agent ICI in UC were based on PD-L1 positivity^51^, and patients with high risk UC after cystectomy and PD-L1 TPS score ≥ 1 appeared to have larger magnitude of benefit to adjuvant nivolumab than the intention to treat population^29^. Consistent with high PD-L1 in SUC, SUC does appear to have a higher objective response rate than pure UC in one study (38% versus 28%)^4^. However, a high TGFb related gene expression score in UC tumors associated with poor response to atezolizumab^52^. One disconnect is how SUC has higher TGFb gene expression yet greater proportion of patients who have response to ICI. The impact of PD-L1 on immune suppression and response to ICI in SUC is an area of active investigation.

In conclusion, this manuscript demonstrated the generation of three variant histologic subtype mouse tumors that initially lacked tumorigenicity, but then acquired growth capacity upon passaging in mice lacking mature B and T cells. Three independent GU oncology pathologists identified BBN966 as SUC, BBN975 as squamous, and BBN964 as a hybrid.

BBN966 had a “double-negative” phenotype that was consistent with a subset of aggressive human SUC. Genes that were upregulated in all three murine tumor cells were found to associate with worse survival in patients with muscle invasive UC whose tumors had high expression of these genes. Human SUC, canine SUC, and BBN966 tumors had increased expression of EMT genes. Consistent with this, BBN966 had low expression of p63 and E-cadherin. BBN966 tumors had high M2 macrophage gene expression, consistent with human SUC. Many SUC tumors are characterized by high PD-L1 IHC staining, and consistent with human SUC, anti-PD1 ICI inhibited growth of BBN966 (but not BBN964) tumors. This work informs our understanding of human SUC, and provides novel variant histology mouse models (including the mouse SUC BBN966) for further study.

## Funding

This study was supported by a Department of Defense Career Development Award (CA220769 to B.A.J.), an NIH NCI R01 (CA235681 to N.M.H.), a Bladder Cancer Advocacy Network Young Investigator Award (to B.A.J.), a Commonwealth Foundation award (to B.A.J. and D.J.M.), the Johns Hopkins Greenberg Bladder Cancer Institute, The John and Camilla Duffy Urologic Oncology Research Fund, and the Johns Hopkins Sidney Kimmel Comprehensive Cancer Center Support Grant NIH NCI P30 (CA006973). A.Y. was supported in part by a Japanese Society for the Promotion of Science Overseas Research Fellowship.

## Conflict of Interest Disclosures

Drs. Hoffman-Censits, Hahn, McConkey, and Johnson hold a pending patent related to the use of an immune cell based biomarker as a prognostic and predictive biomarker in cancer. Dr. Johnson previously served on an Emerging Thought Leaders Advisory Board for Seattle Genetics. Dr. Hahn reports research funding from AstraZeneca, Incyte, Genentech, BMS, Merck, Seattle Genetics, Astex, Principia Biopharma, Pieris, and HTG Molecular Diagnostics; consulting income from Merck, Genentech, GlaxoSmithKline, Ferring, Champions Oncology, Health Advances, Keyquest Health, Guidepoint Global, Seattle Genetics, Incyte, TransMed, CicloMed, Janssen, Pfizer, and Boehringer Ingelheim; and honoraria from Bladder Cancer Academy. Dr. McConkey reports grant support from AstraZeneca, and serves on the advisory boards of Janssen, Rainier, and H3 Biomedicine. No other conflicts of interest were reported.

---

**Supplemental Figure 1.**
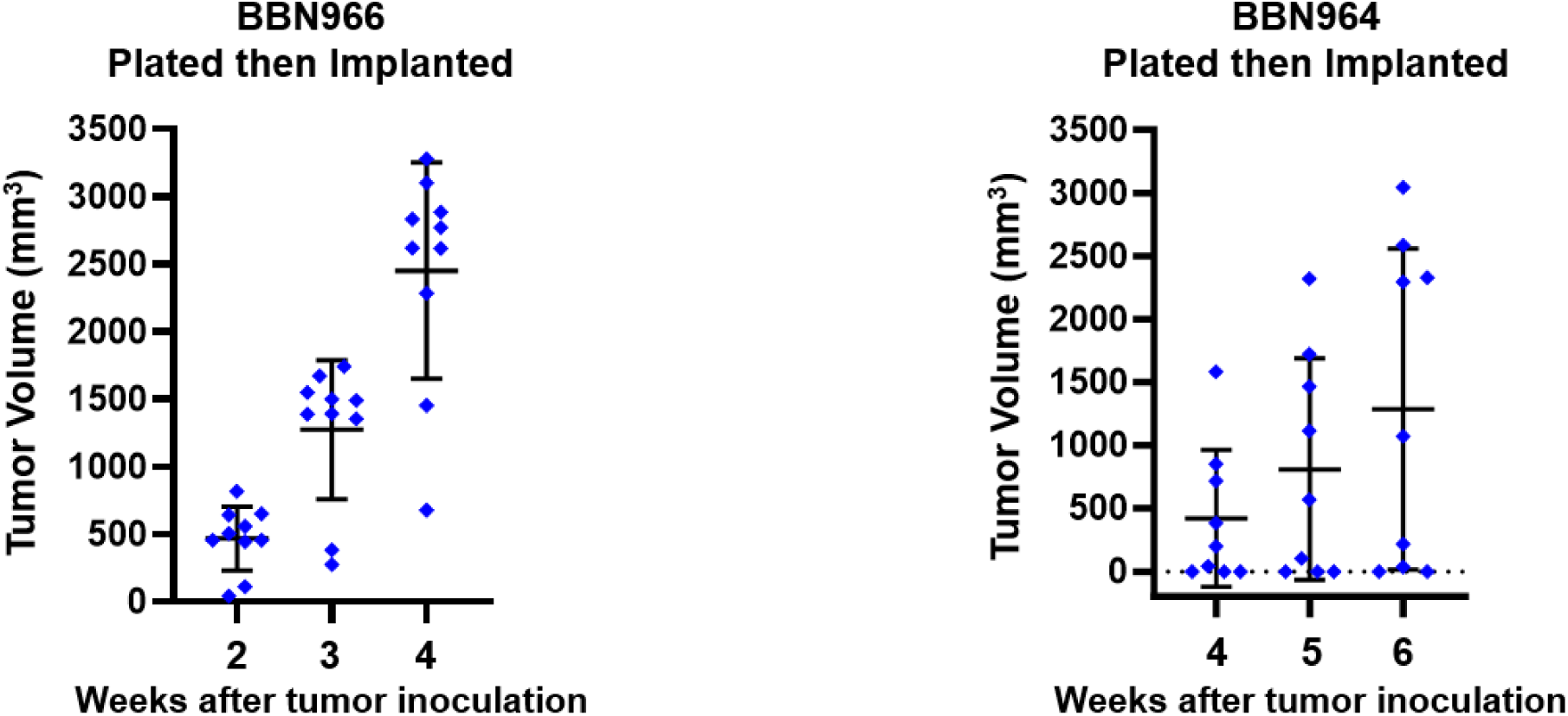
Cells from BBN966 (left) and BBN964 (right) tumors were plated on plastic, then grown to confluence and passaged. Tumor cells were trypsinized and implanted (5x10^6, s/q) into naïve B6 mice.

**Supplemental Figure 2.**
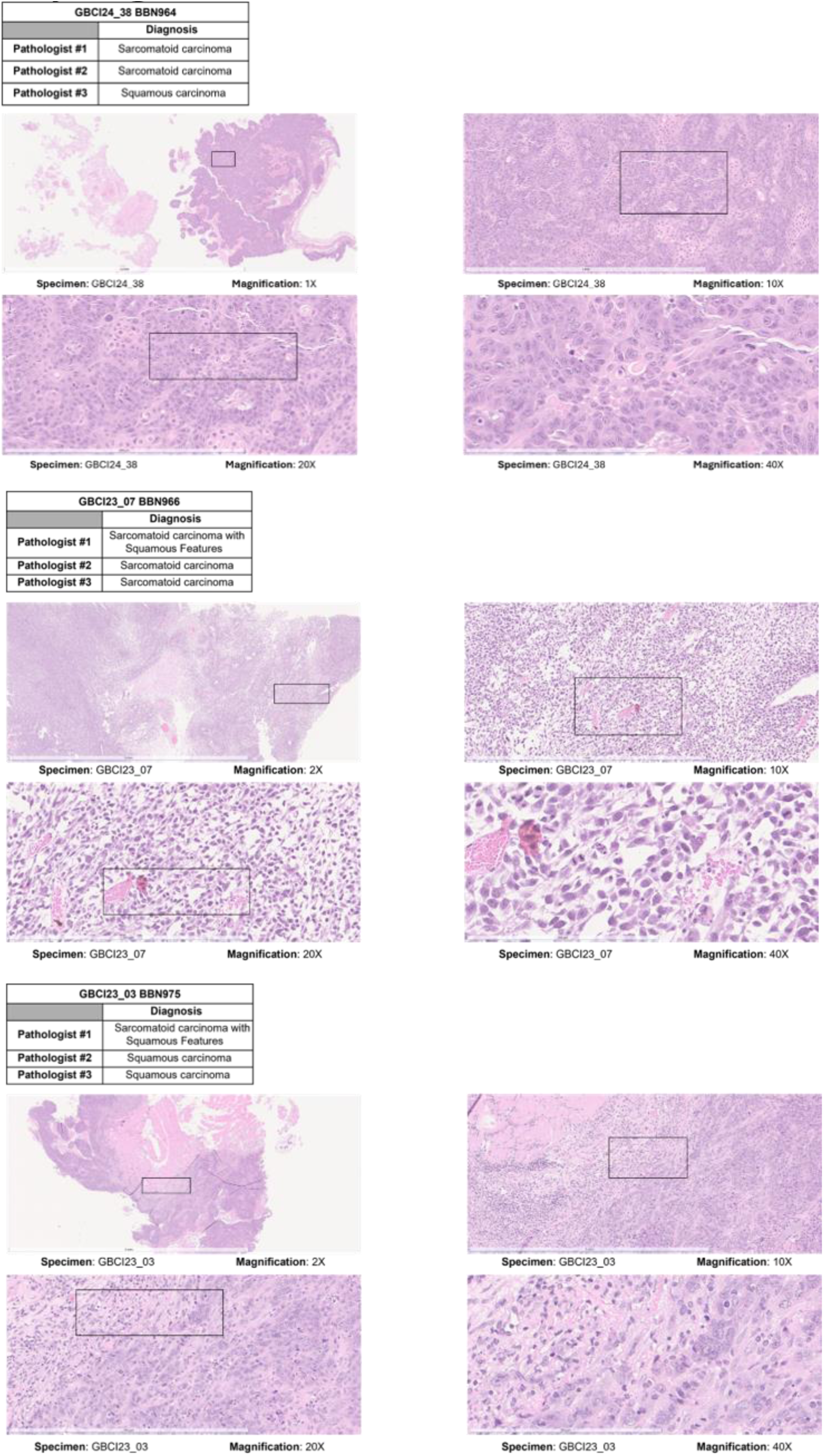
Pictures of BBN964, BBN966, and BBN975 at lower magnifications.

**Supplemental Figure 3.**
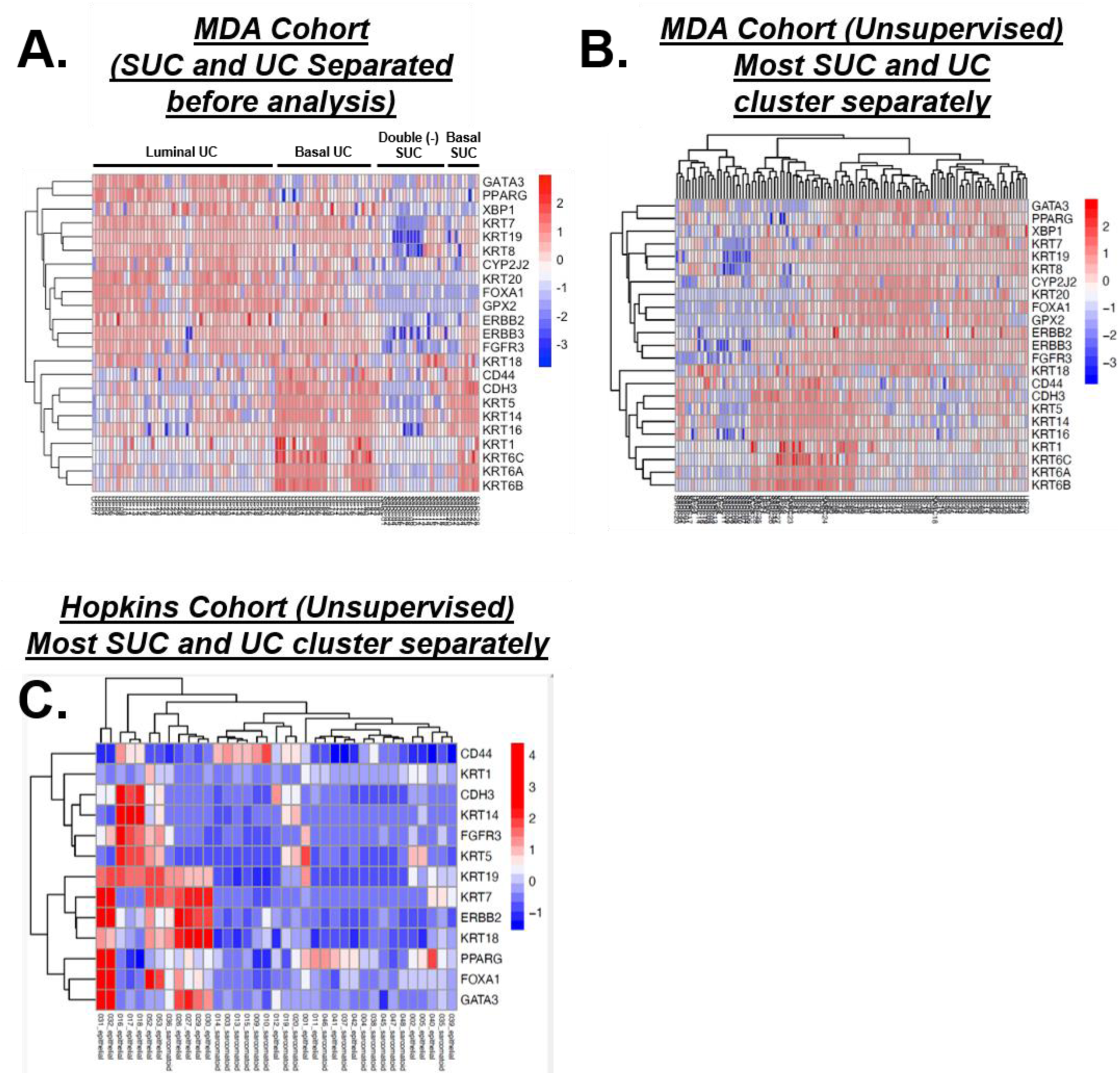
**A.** MD Anderson cohort of unique sarcomatoid and conventional urothelial carcinoma tumors, with sarcomatoid and conventional urothelial carcinoma segregated into separate groups before analysis. **B-C.** Unsupervised clustering of **(B.)** MD Anderson cohort and **(C.)** Hopkins cohort demonstrated that sarcomatoid and conventional urothelial carcinoma generally segregated into unique clusters.

**Supplemental Figure 4.**
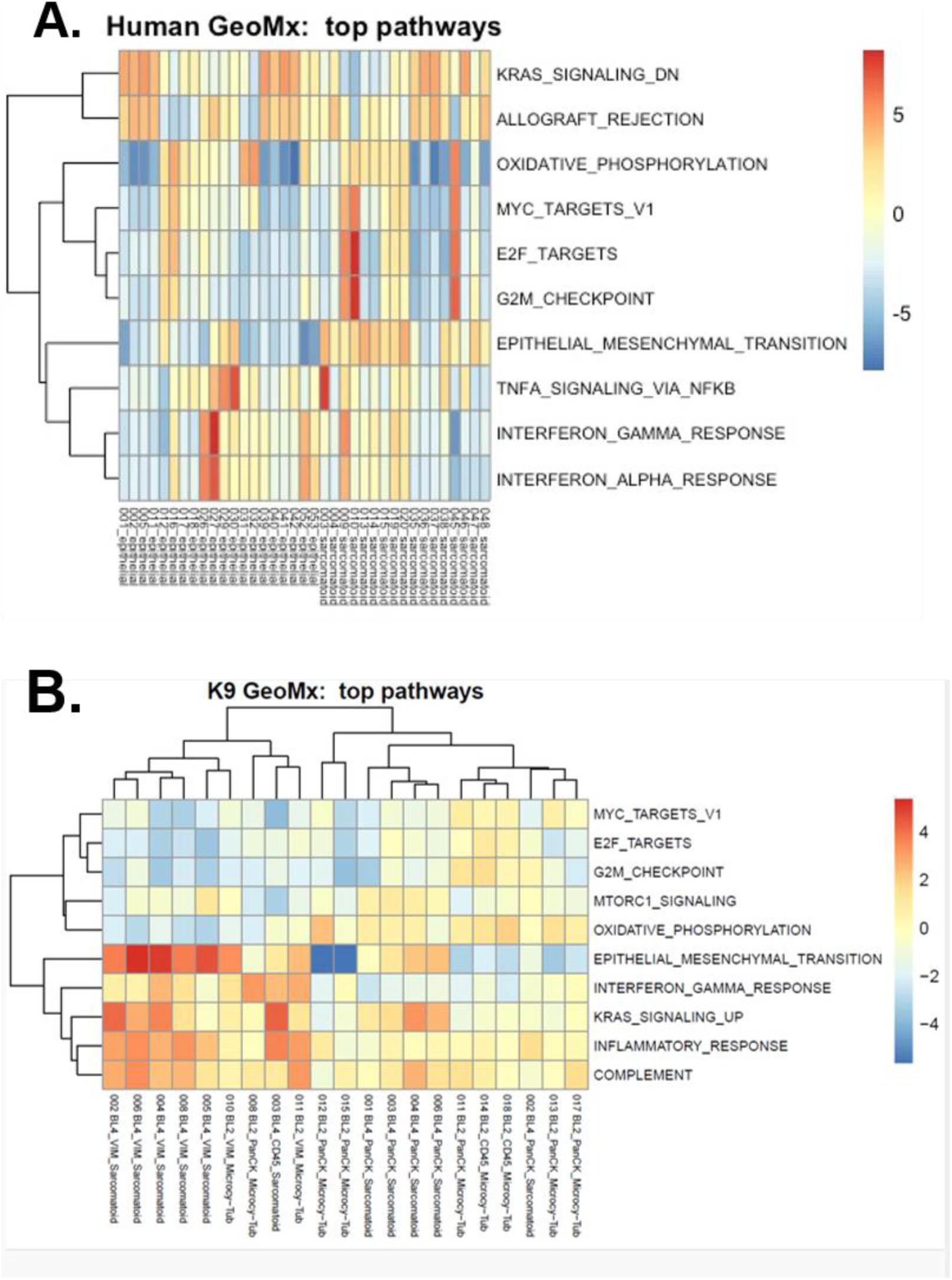
**A.** Pathway analysis generated via spatial transcriptomics analysis of the Hopkins cohort of UC and SUC from multiple ROI of mixed tumors, separated by UC and SUC ROI. **B.** Unsupervised clustering of multiple ROI from a mixed UC and SUC canine tumor, including pathway analysis.

**Supplemental Figure 5.**
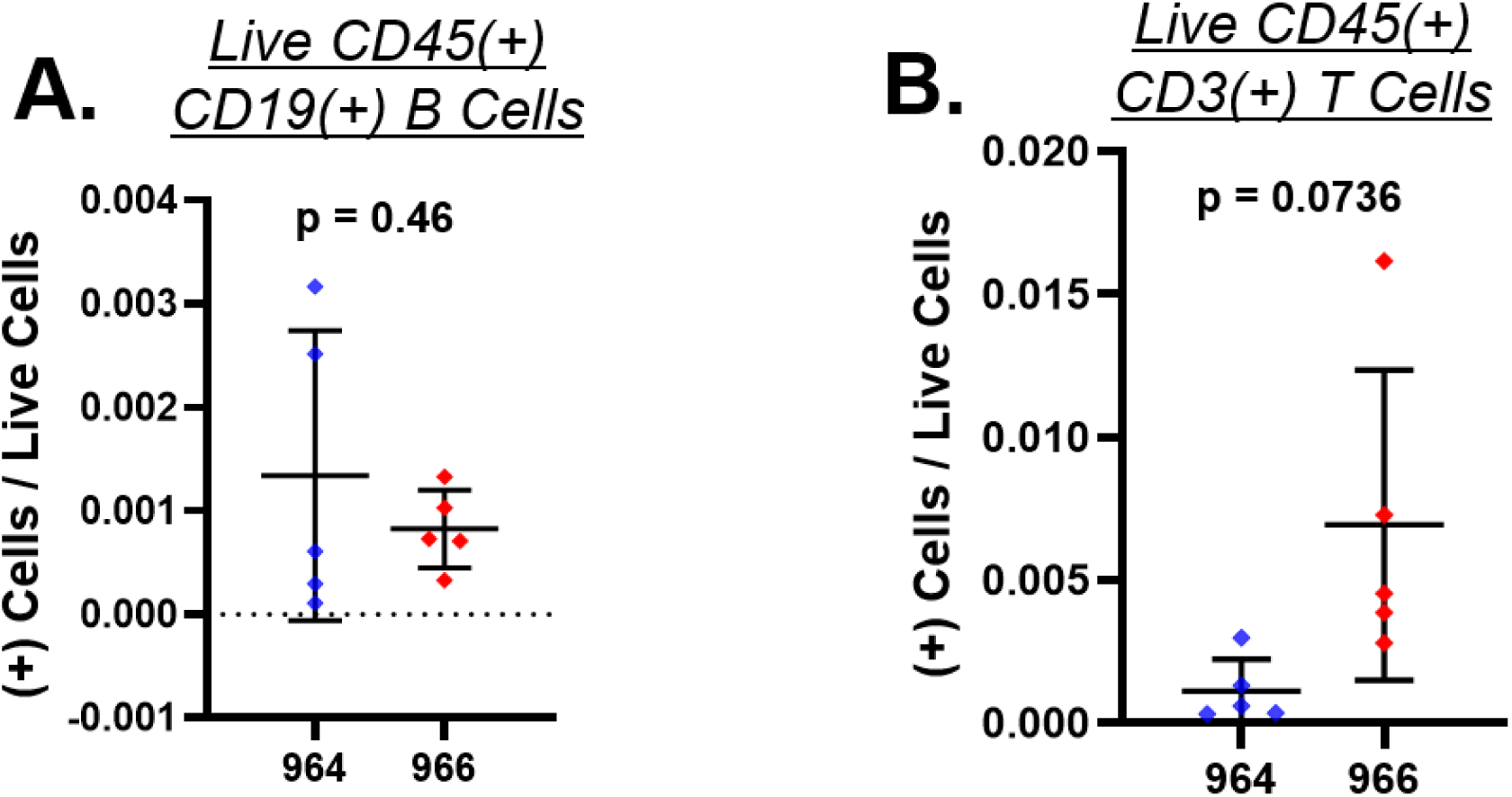
**A-B. (A.)** CD45+CD19+ B cells and **(B.)** CD45+CD3+ T cells as a proportion of total live cells in BBN964 and BBN966 tumors (both passaged in B6 for at least 1 year).

**Supplemental Tables 1-3.**
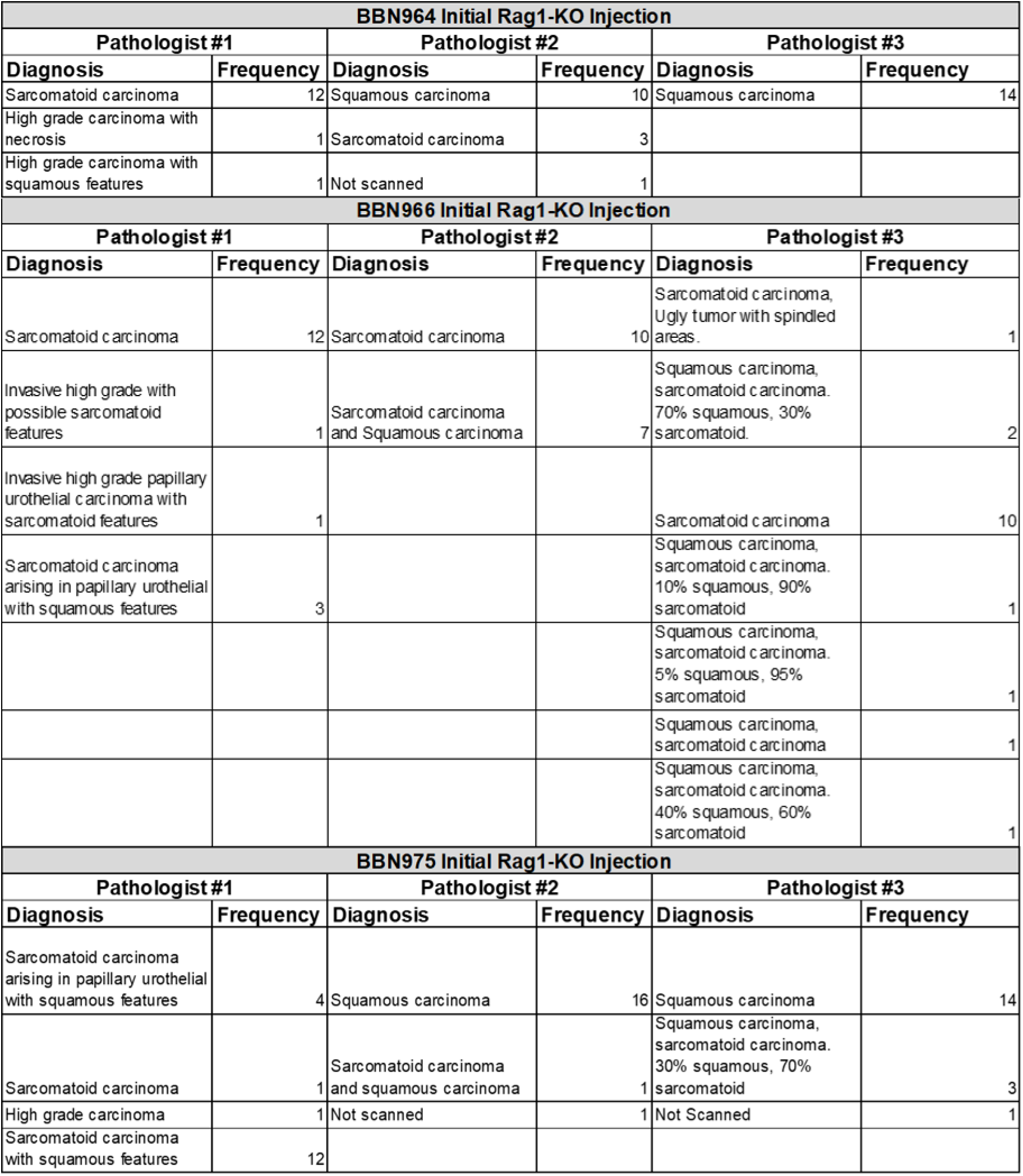
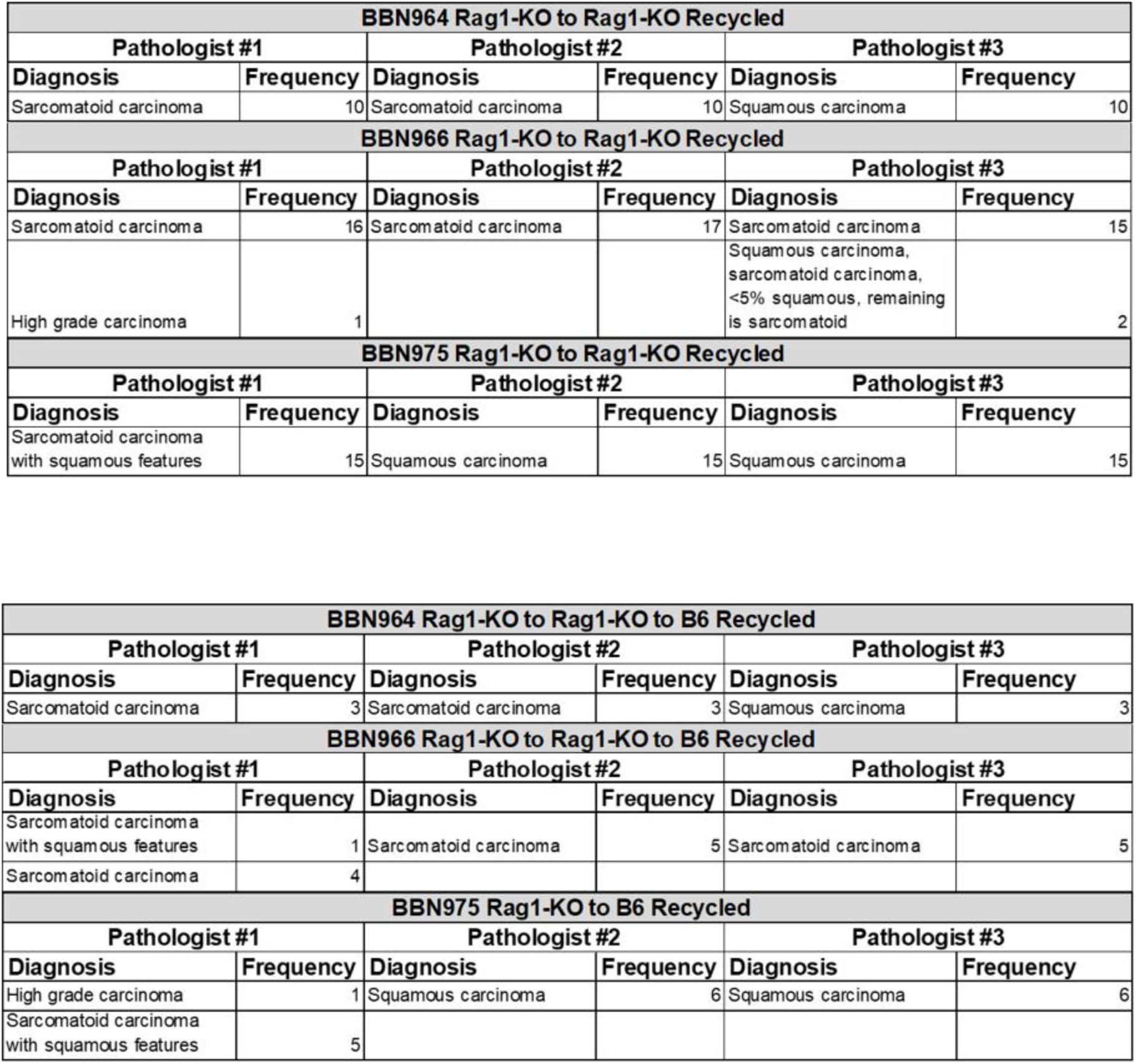
Tables providing additional detail on pathologic diagnoses in **Figure 2**. Specific pathologic diagnoses are provided for tumors from initial Rag1-KO injection (**Sup. Table 1**), tumors from the Rag1-KO to Rag1-KO recycled injection (**Sup. Table 2**), and tumors from the initial B6 implantation following Rag1-KO implantation (**Sup. Table 3**).

